# Biased activation of the vasopressin V2 receptor probed by NMR, paramagnetic ligands, and molecular dynamics simulations

**DOI:** 10.1101/2023.06.06.543947

**Authors:** Gérald Gaibelet, Aurélien Fouillen, Stéphanie Riché, Hélène Orcel, Christiane Mendre, Ali Kanso, Romain Lanotte, Julie Nguyen, Juliette Dimon, Serge Urbach, Rémy Sounier, Sébastien Granier, Dominique Bonnet, Xiaojing Cong, Bernard Mouillac, Hélène Déméné

**Affiliations:** Institut de Génomique Fonctionnelle, Université de Montpellier, CNRS, INSERM, 34094 Montpellier cedex 5, France; Laboratoire d’Innovation Thérapeutique, UMR7200 CNRS, Université de Strasbourg, Institut du Médicament de Strasbourg, 67412 Illkirch-Graffenstaden, (France); Centre de Biologie Structurale, Univ Montpellier, INSERM, CNRS, Montpellier (France)

**Keywords:** GPCR dynamics, Biased Activation, NMR, Molecular Dynamics, Pharmacology

## Abstract

G protein-coupled receptors (GPCRs) control critical intercellular communications by responding to extracellular stimuli and undertaking conformational changes to convey signals to intracellular effectors. We combined NMR, molecular pharmacology, and molecular dynamics (MD) simulations to study the conformational diversity of the vasopressin V2 GPCR subtype (V2R) bound to different types of ligands: the antagonist tolvaptan, the endogenous unbiased agonist arginine-vasopressin, and MCF14, a Gs-protein biased agonist. We developed a double-labeling NMR scheme to study the conformational dynamics: V2R was subjected to lysine ^13^CH_3_ methylation, whereas the agonists were tagged with a paramagnetic probe. Paramagnetic relaxation enhancements were used to validate the ligand binding poses in the MD simulations. We found that the bias for the Gs protein over the β-arrestin pathway involves interactions between the conserved NPxxY motif in the transmembrane helix (TM) 7 and a central hydrophobic patch in TM3, which constrains TM7 and likely inhibits β-arrestin signaling. A similar mechanism was observed for the pathogenic mutation, I130^3.43^N, which constitutively activates the Gs protein without concomitant β-arrestin recruitment. This mechanism resembles to opioid receptors findings indicating common patterns in class A GPCRs.

## Introduction

GPCRs are the largest protein family in the human genome and key players in cell signaling. As transmembrane receptors, they sense extracellular stimuli and trigger signal transduction cascades inside the cell, principally via G protein and β-arrestin (βArr) pathways.^[1]^ GPCR ligands may preferentially modulate some signaling pathways over others. This functional selectivity is known as GPCR ligand bias,^[2]^ which enables better control of therapeutic effects, opening new avenues to drug discovery. GPCR-biased signaling may even occur in the *apo* state, in certain constitutively active (often pathological) mutants.^[3,4]^

Both X-ray crystallography and cryo-electron microscopy (cryo-EM) revealed remarkably unified structural rearrangements upon class A GPCR activation stabilized by G proteins or βArrs, characterized by outward displacement of the TM6 and inward movement of TM7.^[5–9]^ However, receptor conformational changes before coupling with intracellular partners, essential for biased signaling, are highly dynamic and poorly understood. While spectroscopies in solution such as NMR or fluorescence and MD simulations have provided valuable information,^[10–17]^, the bridge with static structures is still missing.^[14,18–20]^

V2R regulates the renal antidiuretic response in mammals.^[21]^ Binding of arginine vasopressin (AVP) to V2R activates both Gs protein and βArr pathways associated to urine concentration,^[21]^ and cell growth and proliferation, respectively.^[22,23]^ V2R is a major therapeutic target for water balance disorders.^[24]^ Loss-of-function mutations result in the congenital nephrogenic diabetes insipidus (cNDI),^[25]^ characterized by excessive urine voiding and dehydration^[26]^ The mutant receptors are usually misfolded and retained in the endoplasmic reticulum.^[27]^ A class of benzazepine compounds, the MCF series, has been found to rescue expression and signaling of some cNDI mutants. They display promising biased profiles of long-lasting activation of the Gs protein pathway, without triggering βArr-mediated internalization.^[28]^ Some pathological gain-of-function V2R mutations lead to the nephrogenic syndrome of inappropriate antidiuresis (NSIAD), characterized by hyponatremia.^[29]^ Several of these mutants, such as I130^3.43^N (superscript refers to Ballesteros-Weinstein nomenclature)^[30]^, display constitutive activities only for Gs protein, mimicking GPCR-biased signaling.^[31–33]^ High-resolution V2R structures have been recently obtained by us and others, only of active states in complex with AVP and either Gs ^[34–36]^ or βArr1 proteins.^[37]^ No structure of V2R in an inactive or biased state is yet available. Hence, little is known about V2R ligand or mutation-induced bias and dynamics, apart from our preliminary fluorescence study showing that AVP and the Gs-biased agonist MCF14 induced distinct V2R conformations on the intracellular side of TM7 and helix 8 (H8).^[13]^ Here, we combine functional cell pharmacology, NMR spectroscopy, and enhanced-sampling MD simulations to identify comprehensively V2R conformations associated with the Gs-biased agonist MCF14 and the constitutively Gs-active mutant I130^3.43^N in comparison with unbiased agonist AVP and the antagonist tolvaptan (TVP). To assess the ligand binding poses, we developed a double-labeling scheme using lysine ^13^CH3 methylation on V2R and paramagnetic tagging of the ligands. The Paramagnetic Relaxation Enhancement (PRE) data were in agreement with the measures from MD simulations. We also could identify distinct receptor conformations associated with the Gs-biased pathways, pointing out the role of a central hydrophobic patch in TM3 and the pivotal NP^7.50^xxY motif in TM7 in biased V2R activation mechanism.

## Results and Discussion

### NMR Sensors and Paramagnetic Ligands

To characterize the conformational changes of V2R, we introduced NMR sensors by methylating its lysines with two ^13^CH3 groups for heteronuclear ^1^H-^13^C NMR.^[38,39]^ The V2R construct has three lysine, K100^2.65^, K116^3.29^ and K268^6.32^ (Figures 1A, S1A and S2). K100^2.65^ lies on the rim of the orthosteric pocket, whereas K116^3.29^ is located within the pocket. K268^6.32^ belongs to the cytoplasmic extremity of TM6, which makes it an appropriate sensor of canonical activation, considering the pivotal role of TM6 therein.[8,9],[35–37]

**Figure 1.**
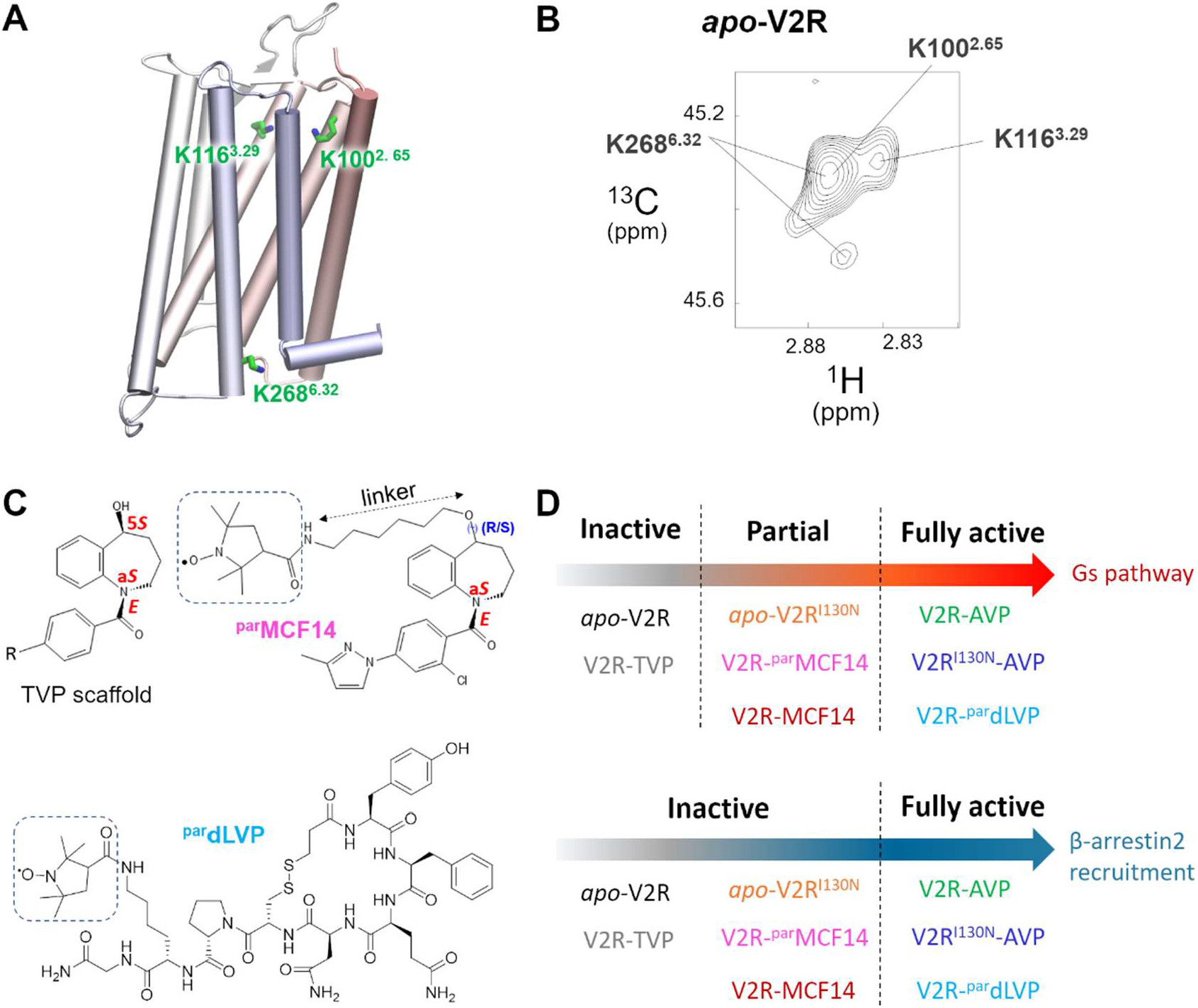
NMR probes, paramagnetic labels, and functional profiles of the systems studied. (A) Position of the three lysine sensors in inactive V2R. (B) ^1^H-^13^C HMQC NMR spectrum of ^13^CH3-Lys-V2R. (C) Stereochemical configuration of TVP-type V2R ligands and chemical structures of the paramagnetic agonists. The paramagnetic tags are boxed. The chiral center of ^par^MCF14 is marked. (D) Functional profile of systems studied in the MD simulations (see also Table 2).

Peak assignment of the heteronuclear ^1^H-^13^C HMQC spectrum of methylated V2R (Figure 1B) was obtained by site-directed mutagenesis of lysine into arginine (Figure S3). Contrary to K100^2.65^ and K116^3.29^, which have only one correlation peak, K268^6.32^ is represented by a major broad peak and at least one minor peak, indicating conformational heterogeneity at the cytoplasmic end of TM6. NMR and mass spectrometry (MS) data eliminated the possibility of incomplete methylation labeling of lysine residues that may result into several resonances of the K268^6.32^ signal (Figure S2).

Accurate prediction of the ligand binding pose is critical for MD simulations. Since *i*) only the AVP binding pose is known from the cryo-EM structures, and *ii*) TVP and MCF14 structurally differ from AVP (Figure S1B), we used ligand paramagnetic tagging through covalent addition of a TEMPOL group to verify the ligand binding poses predicted by MD simulations. The TEMPOL group, containing a nitroxide probe, was linked to either MCF14 or to the deaminated lysine-vasopressin (dLVP) to obtain their paramagnetic forms (^par^MCF14 and ^par^dLVP, respectively) (Figures 1C and S1B). The dLVP, an AVP analog that behaves as AVP towards V2R antidiuretic activity,^[40]^ was used to facilitate chemical synthesis (Scheme S1) and to validate the paramagnetic approach. PREs of the ^1^H nuclei, a well-known source of distance information, provide the distance estimation of TEMPOL to the methyl groups of K100^2.65^ and K116^3.29^, which can be compared directly with the MD simulations. The paramagnetic tag position in ^par^MCF14 was determined based on the presumed binding pose of TVP-like V2R ligands. Since they bind V2R mostly in *E*-a*S-*5(*R*/*S*) configurations, ^[41]^ we chose the C5 position to insert a linker (Fig. 1C), in order to orient the paramagnetic nitroxide probe outward from the pocket. Introduction of the paramagnetic tag revealed no alteration in the signaling profiles of both ligands, although their affinity and potency were reduced (Figure 2A and Table 1). ^par^dLVP remains a fully unbiased agonist, whereas ^par^MCF14 is a Gs-biased partial agonist like its parent compound (Figure 1D and Table 2).

**Table 1:** Ligand affinity and potency in living cells^[a]^

|  | K<sub>i</sub> (nM)<sup>[b]</sup> | EC<sub>50</sub> (nM)<sup>[b]</sup> |  |
| --- | --- | --- | --- |
| | | cAMP | βArr2 |
| AVP | 0.93 ± 0.3 (n=5) | 0.11 ± 0.01 | 0.85 ± 0.10 |
| <sup>par</sup>dLVP | 273.5 ± 39.7 | 37.5 ± 3.2 | 110.5 ± 18 |
| MCF14 | 7.9 ± 2.3 | 40.9 ± 11.6 | n.a.<sup>[c]</sup> |
| <sup>par</sup>MCF14 | 169.7 ± 24.7 | 647.6 ± 37 (n=5) | n.a. |

[a] Measured in cells expressing the V2R construct used for NMR (Figure S1A).
[b] Values are mean ± SEM (n = 3, unless otherwise indicated)
[c] n.a.= not applicable.

**Table 2:**
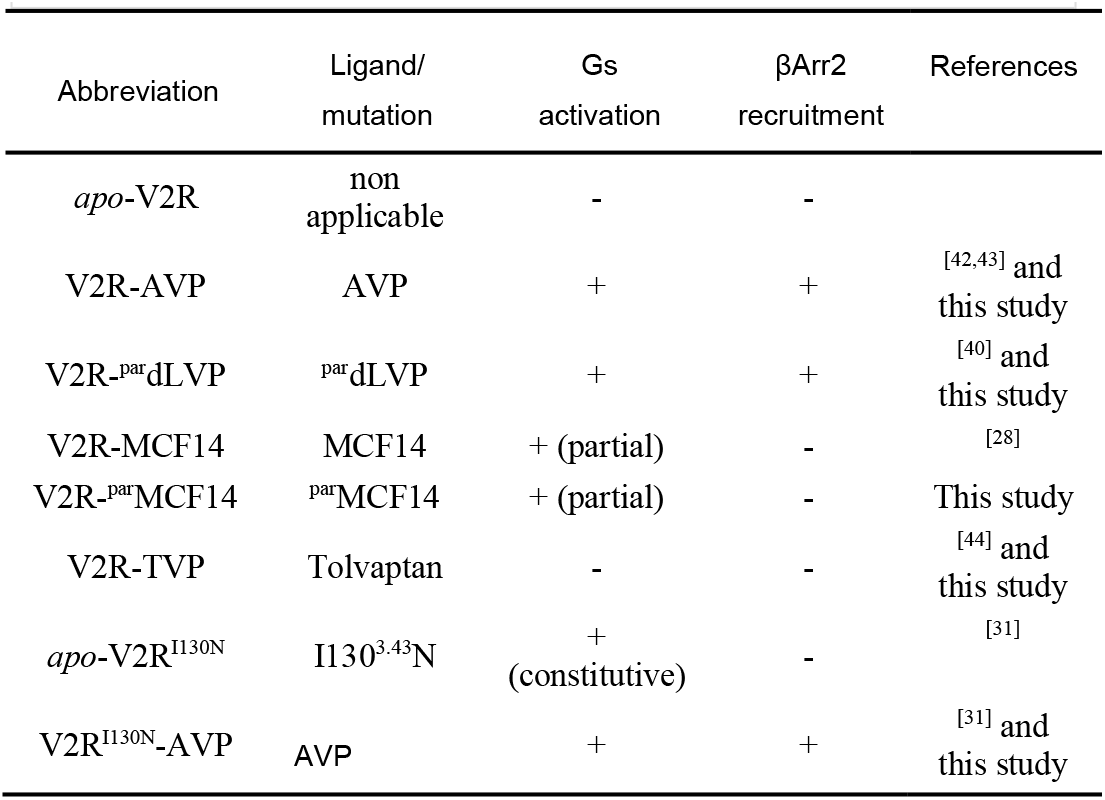
V2R systems studied

**Figure 2.**
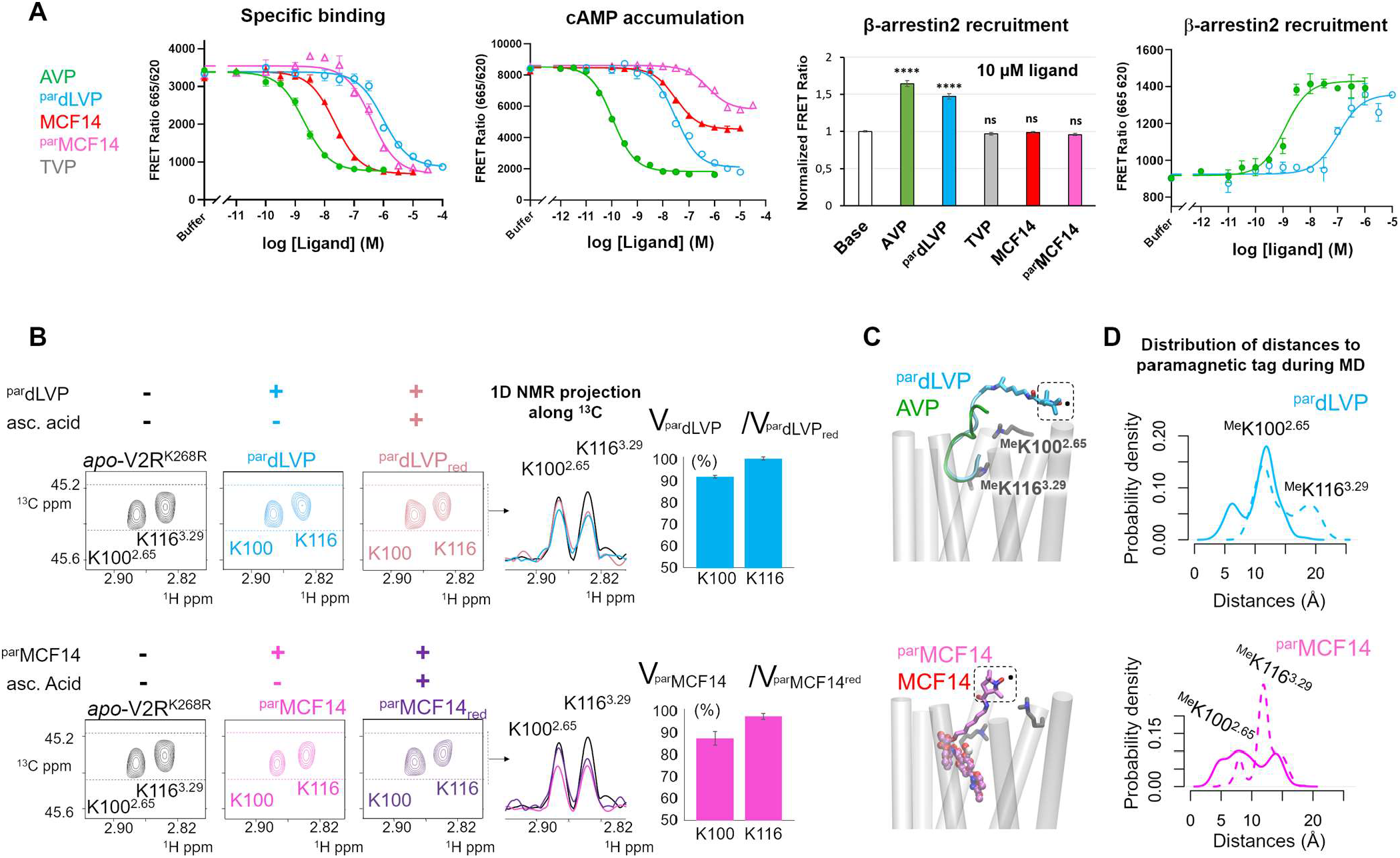
Pharmacological properties and binding poses of the ligands in V2R. (A) Binding properties (left), effects on cAMP accumulation (central-left) and β-arrestin2 recruitment (center-right and right). TR-FRET was expressed as FRET ratio (665 nm/620 nm x 10,000). ****, p<0.0001; ns, not significant (p>0.05). (B) Extracted HMQC spectra of K100^2.65^ and K116^3.29^ resonances of V2R K268^6.32^R in *apo* form (black), bound to ^par^dLVP (sky blue) or ^par^MCF14 (magenta), and reduced by ascorbic acid (light violet and mauve taupe). 1D spectra (right) correspond to 1D projections along the ^13^C dimension of ^1^H rows between dashed lines. Ratios of peak volumes in the paramagnetic and diamagnetic (ligand_red_) forms are shown to the right. (C) Representative positions of tagged and non-tagged agonists during MD simulations. The nitroxyde cage is squared. (D) Probability density distribution of the distances between the TEMPOL nitroxide ion and the methyl groups of K100^2.65^ (plain line) and K116^3.32^ (dashed line) during the MD simulations.

### Ligand Binding Poses by NMR and MD

The PREs were estimated by comparing the volumes of the V2R lysine correlation peaks in complex with the tagged ligands, in the paramagnetic and diamagnetic states (Figure 2B). We used V2R^K268R^ for this purpose, to simplify the NMR spectra. The K268^6.32^R mutation had negligible impacts on the ligand affinities and PREs compared to the wild-type (wt) receptor (Table S1 and Figures 2A and S4). Addition of the paramagnetic ligands induced a decrease in the peak volumes of K100^2.65^ and K116^3.29^ (Figure 2B), due to the distance-dependent relaxation effects of the unpaired ion in TEMPOL and the ligand-induced conformational changes. Addition of 5 equivalents of ascorbic acid as a reducing agent attenuated the paramagnetic interactions, resulting in an increase in the lysine peak volumes.

This effect was more remarkable on K100^2.65^ than K116^3.29^ for both ^par^dLVP and ^par^MCF14, suggesting that the side-chain methyls of K100^2.65^ are closer to the paramagnetic center than that of K116^3.29^ (Figure 2B). Of note, ^par^dLVP maintained the same binding pose in the MD simulations as AVP in the cryo-EM structures (Figure S5A).^[35–37]^ The paramagnetic tag swayed at the entrance of the orthosteric pocket (Figure 2C). The paramagnetic center in ^par^dLVP was on average closer to the K100^2.65^ methyl groups than those of K116^3.29^ (Figures 2D and S6), in line with the NMR measurements. In the case of ^par^MCF14, the ligand used in the NMR studies was a mixture of two enantiomers (Figure 1C) and we examined both of them in the MD simulations. R-^par^MCF14 turned out to be highly mobile in the pocket during the simulations (Figure S5A), indicating low affinity. This is consistent with the fact that V2R favors the *S*-configuration of TVP-type ligands. S-^par^MCF14 was stable in the pocket and adopted the same binding pose as the untagged MCF14, suggesting that this enantiomer was the active one. Therefore, R-^par^MCF14 was discarded and hereafter, we refer to S-^par^MCF14 as ^par^MCF14.

The paramagnetic center in ^par^MCF14 was closer to the K100^2.65^ methyl groups than those of K116^3.29^, in line with the NMR results (Figures 2D and S6). The paramagnetic ligands were slightly more mobile than their untagged counterparts during the simulations (Figure S5A), in agreement with their lower affinity in the cell assays. Therefore, the MD simulations were in agreement with NMR and pharmacological data.

Since TVP analogs could bind V2R in both *E*-a*S-*5*R* and *E*-a*S-*5*S* configurations, preferably in *E*-a*S-*5*S*,^[41]^ we tested both enantiomers of TVP (*E*-5*R*-TVP and *E*-5*S*-TVP, Figure S1) by MD simulations. In addition, *endo*/*exo* isomers of TVP and MCF14 were also tested (Figure S1). We found that the *endo-*isomer was the most stable in the V2R pocket. Both enantiomers of TVP turned out to bind stably in the V2R pocket, in slightly different binding poses (Figure 3A and Figure S5). MCF14 exhibited a similar binding pose to TVP, which was expected given their chemical similarity (Figure 3A).

**Figure 3.**
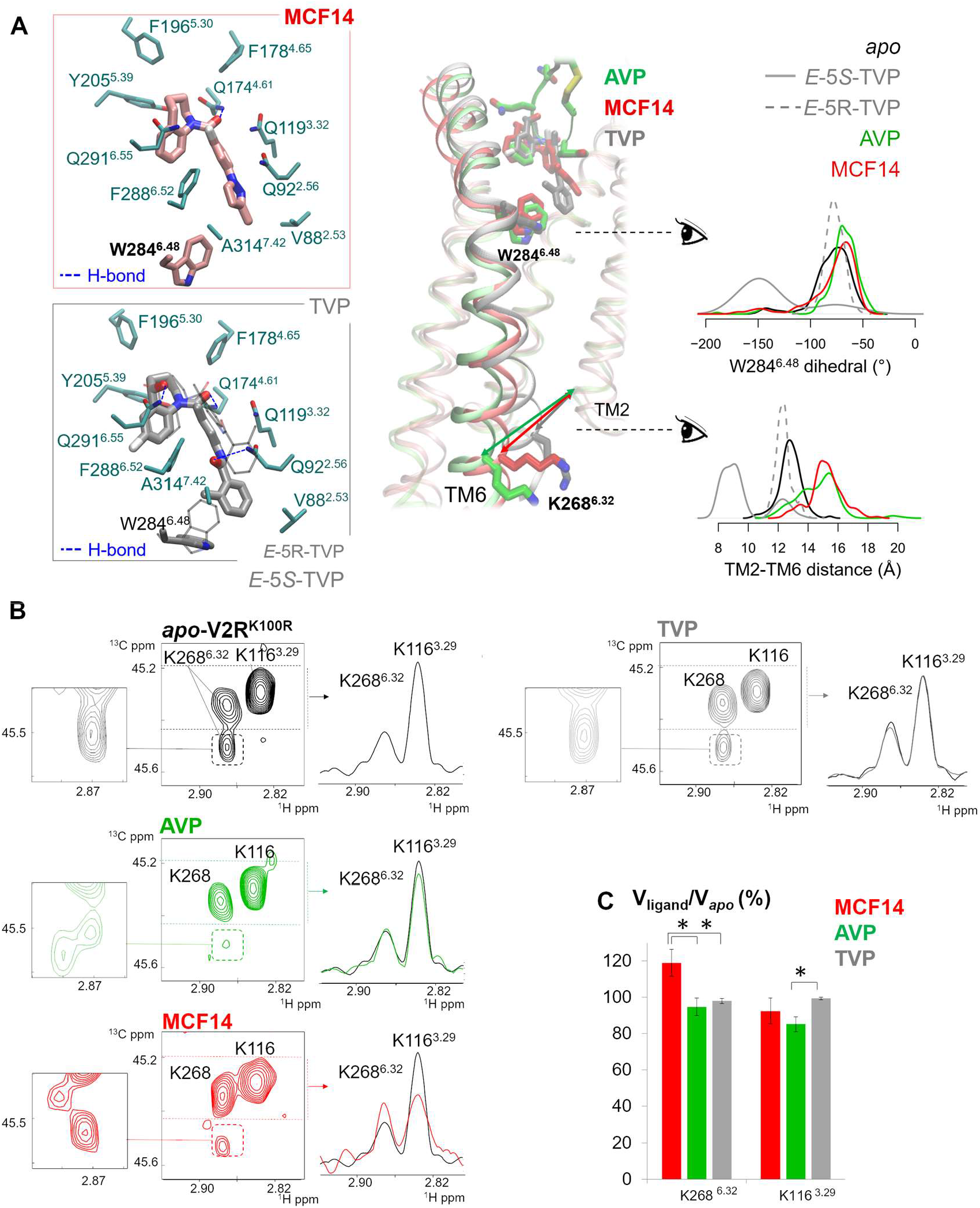
Ligand-induced V2R conformational changes in TM6. (A) Ligand binding poses, W284^6.48^ side-chain orientation and TM6 position observed in MD simulations. The binding poses of *E*-5*S*-TVP and *E*-5*R*-TVP are shown in sticks and lines, respectively. TM2-TM6 distance was measured by the center-of-mass distance between the backbones of I74^2.39^-F77^2.42^ and V266^6.30^-T269^6.33^. Right panel shows the density distribution of W284^6.48^ sidechain dihedral angle *χ*_2_ and the TM2-TM6 distance during the MD simulations. (B) HMQC spectra of the K100^2.65^R mutant in *apo* form (black) and in the presence of an excess of AVP (green), MCF14 (red) and TVP (gray). 1D spectra on the right correspond to 1D projections along the ^13^C dimension between the two dashed lines. For clarity, minor (zoom on the left) and major peaks are shown at two different level of noise. Spectra represented in (B) were recorded on samples originating from the same batch of V2R purification, except for TVP, whose reference spectrum is shown in Figure S10. **(C)** Analysis of peak volume changes of the major peaks upon ligand binding. Values are mean ± SD of three (TVP and AVP) or four (MCF14) technical replicates, where statistical significance is assessed by T-test (*, p < 0.05).

### Correlation of V2R Conformational Changes with Ligand and Mutation Signaling Profiles

Having determined the ligand binding poses, we analyzed the structural features associated with biased and unbiased V2R activation by solution NMR and MD simulation. Among the eight simulations systems studied (Figure 1D and Table 2), we included the I130^3.43^N mutant, which was previously described to constitutively activate the Gs pathway,^[31]^ consistent with our assay outcome (Figure S7). Upon AVP stimulation, the mutant could activate both Gs and βArr pathways, indicating that AVP overrides the mutational effect on the signaling preference (Figure S7). Therefore, the eight systems divide into three categories: inactive, Gs activation, and unbiased activation (Figure 1D and Table 2).

All the six systems in the activation category showed apparent TM6 outward movements with respect to TM2 on the cytoplasmic side (Figures 3A, S5B, S8, and S9). V2R-AVP exhibited the highest flexibility and distinct conformational clusters at the intracellular end of TM6.Interestingly, *E*-5*R*-TVP stabilized TM6 in the initial position, like *apo*-V2R, whereas *E*-5*S*-TVP induced further closure of TM6. Moreover, *E*-5*S*-TVP formed additional H-bonds with V2R and induced distinct orientations of W284^6.48^ side chain contrasting all the other seven systems (Figures 3A and S8A). These suggest higher affinity and efficacy of *E*-5*S*-TVP than *E*-5*R*-TVP for V2R, consistent with the previous study of TVP analogs. ^[41]^.

W284^6.48^ is part of the conserved CW^6.48^xP motif in class A GPCRs, known as the “toggle switch” of receptor activation.^7^

Therefore, TVP likely antagonizes V2R by blocking this activation switch through its *E*-5*S* configuration.

The TM6 movements observed in the MD simulations were in line with the changes in the NMR correlation peaks of K268^6.32^. To observe clear signals, we used the V2R^K100R^ mutant that preserves the signaling properties of wt V2R (Figure S10 and Table S2). In its *apo* form, K268^6.32^ showed at least two resonance peaks, the main peak is also large and weak, indicating conformational heterogeneity at TM6 intracellular end (Figure 3B). Binding of the antagonist TVP caused no significant change.. By contrast, binding to AVP resulted in a narrowing of the K116^3.29^ and K268^6.32^ peaks, whereas the K268^6.32^ minor peak (at 2.86/45.2 ppm) was split into two components of equal but weaker intensities. In addition, MCF14 caused an increase in the line widths of the K268^6.32^ and K116^3.29^ major peaks, together with an increase in the peak intensity of K268^6.32^ (Figure 3B-C). As with AVP, the smallest peak of K268^6.32^ was also split into two peaks of unequal intensities but at different chemical shifts from AVP, one being close to the small peak found in the inactive states (*apo* and TVP-bound). Hence, the NMR spectra of V2R in its agonist-bound forms indicate at least three different conformations of TM6. The cryo-EM maps of V2R complexed with AVP and Gs showed as well three conformations, suggesting that they may vary in Gs protein engagement and in the capacities of nucleotide exchange, as reported for the adenosine A2A receptor.^[45]^Altogether, the MD and NMR data show that TVP, MCF14, and AVP differently impact TM6 intracellular extremity.

### Biased Activation Mechanism

MD simulations showed distinct TM7-H8 conformations in V2R-MCF14 compared to V2R-AVP (Figure 4). MCF14 bound deeper in the orthosteric pocket and inserted between TM2 and TM7, altering the conformations of the conserved Na^+^-binding site. The NP^7.50^xxY motif moved inward and Y325^7.53^ formed water-mediated or direct H-bonds with T134^3.47^ (TM3). H8 exhibited more constrained conformations and was closer to TM1 than in V2R-AVP (Figure 5A). ^par^MCF14 gave rise to the same phenomenon, despite larger fluctuations in the orthosteric pocket due to lower affinity (Figure S6A and S9).

**Figure 4.**
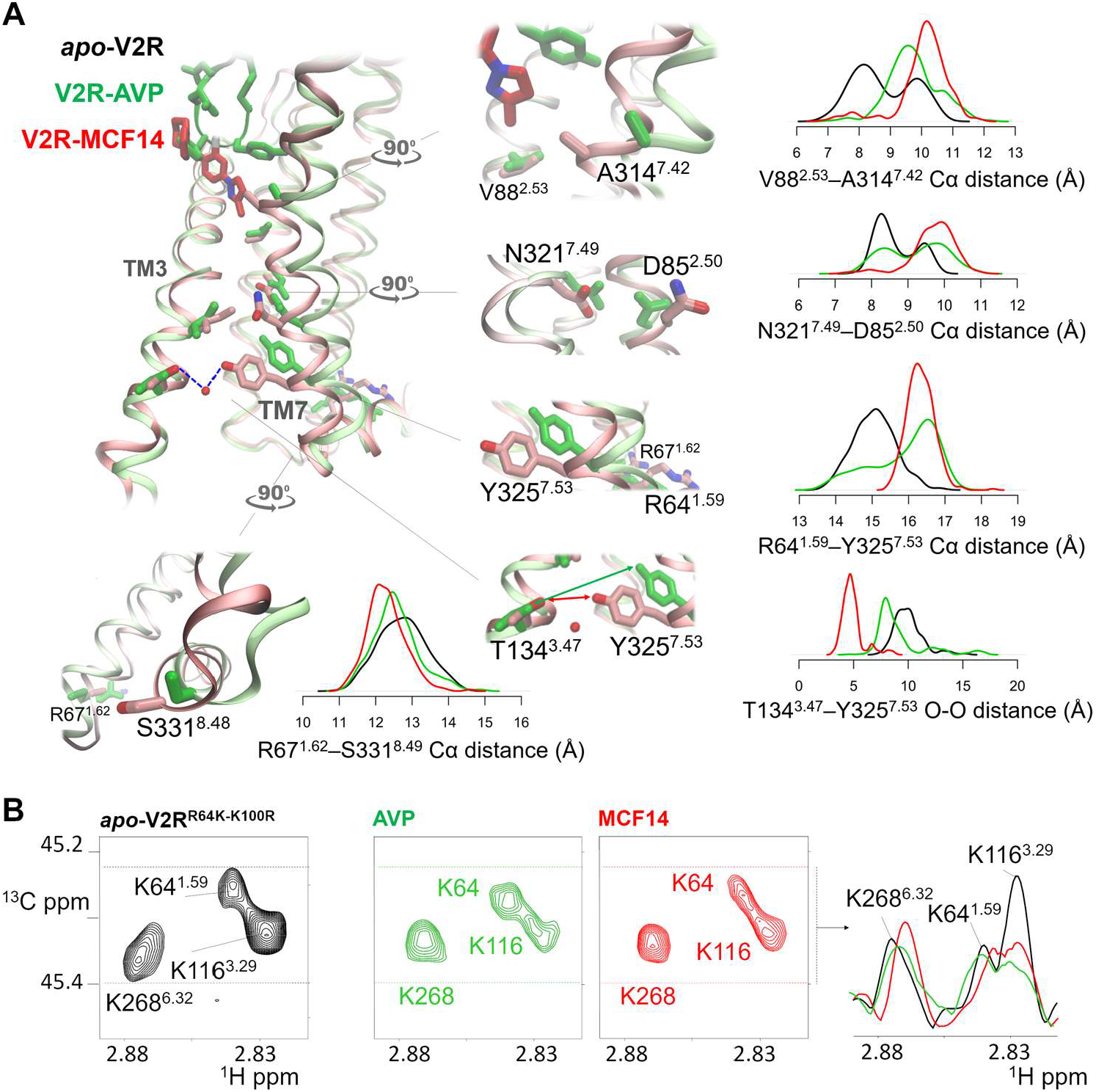
Proposed mechanism of G protein bias by MCF14. (A) Insertion of MCF14 between TM2 and TM7, and successive conformational changes leading to H8 and TM1 compaction. (B) Spectral signature of the ligand nature on the K64^1.59^ NMR probe

**Figure 5.**
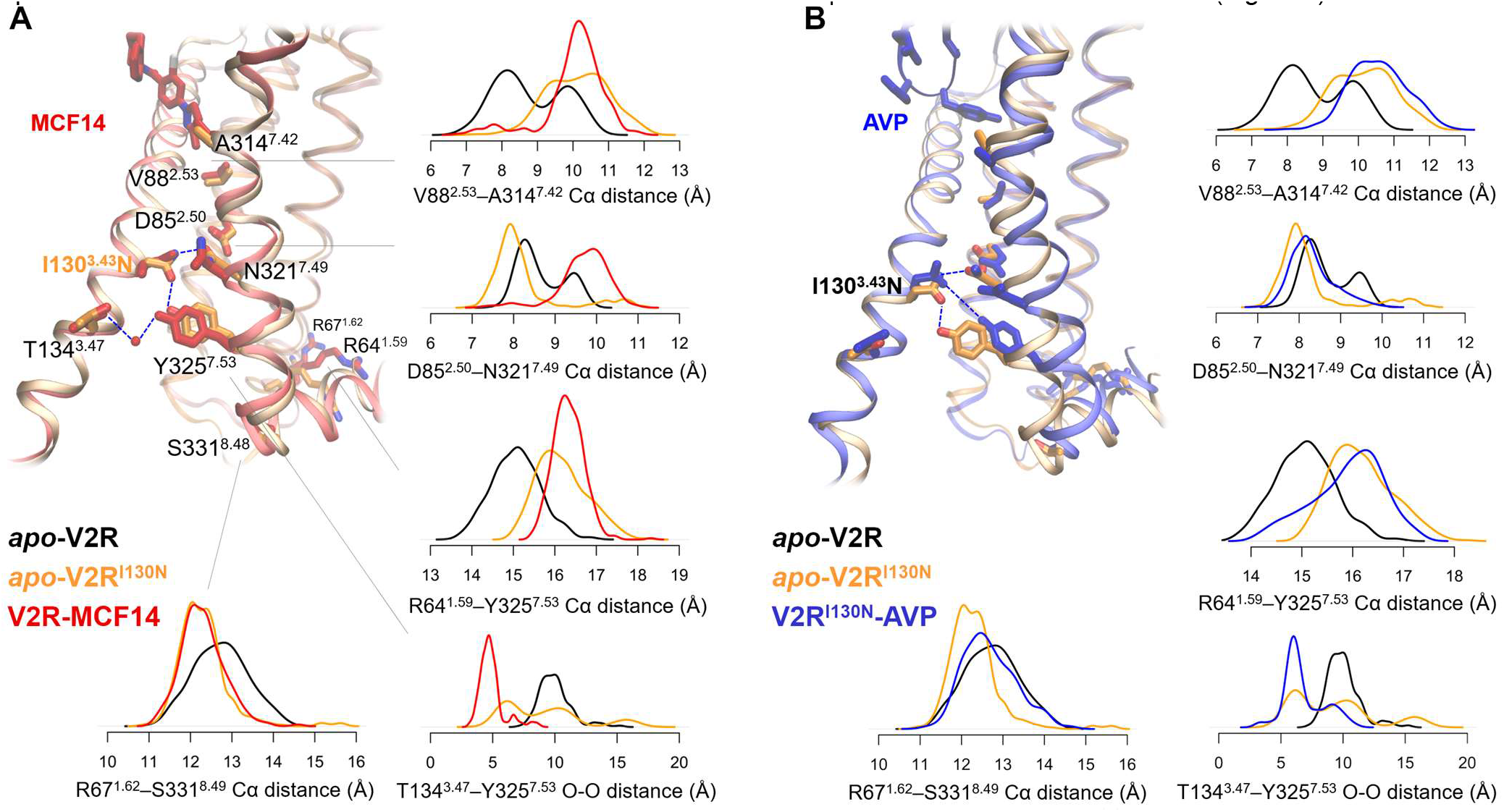
Conformational changes of TM7 and H8 induced by the mutation I130^3.43^N. (A) Changes observed in the specific Gs-signaling pathway promoted by MCF14 or I130N mutation. (B) Allosteric override of the Gs-signaling preference by AVP binding to V2R^I130N^

Remarkably, *apo*-V2R^I130N^ showed similar TM7-H8-TM1 conformations to those in V2R-MCF14 on the intracellular side. These conformations were stabilized as well by a direct H-bond between Y325^7.53^ of the NP^7.50^xxY motif and the mutated I130^3.43^N residue in TM3. The H8-TM1 region was also more compact in V2R-MCF14 and *apo*-V2R^I130N^ (Figures 5A and S9B). Interestingly, stimulation of V2R^I130N^ by AVP (AVP-V2R^I130N^) recovered the wt V2R-AVP-like conformations at H8-TM1 (Figure 5B). Since the I130^3.43^N variant could recruit βArr2 upon AVP stimulation (Figure S7), the distinct H8-TM1 conformations in V2R-MCF14 and *apo*-V2R^I130N^ suggest a link to their lack of detectable βArr signaling, clearly differentiable from V2R-AVP and V2R^I130N^-AVP.

To test this model, we endowed V2R with a lysine probe in TM1, by mutating R64^1.59^ to lysine. This position was chosen, as the sole harboring of an arginine and close to TM7. To minimize spectral crowding, we introduced the R64^1.59^K mutation in Overall, the present study is in line with our previous findings using fluorescent probes in V2R TM6 and TM7-H8 junctions,^[13]^ but the present work provides a far more comprehensive mechanistic view. We elicit here a mechanism where distinct conformations of TM7 lead to the compactness of H8 toward TM1, likely associated with G protein bias (Figure 6). The biased agonist MCF14 triggered these conformational changes allosterically, through the conserved Na^+^-binding site in TM2 and the NP^7.50^xxY motif in TM7. The I130^3.43^N mutation, however, acted directly on the NP^7.50^xxY motif and resulted in similar H8-TM1 conformations to those in V2R-MCF14. Similar patterns at H8 associated with Gi protein bias have been observed on the μ-and κ-opioid receptor. ^[14,46]^

**Figure 6.**
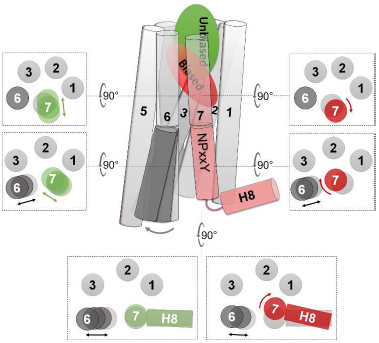
Proposed scheme of V2R conformational changes induced by G protein-biased and unbiased agonists, viewed perpendicularly and from the extracellular side.

Residue I130^3.43^ may play a particular role in GPCR activation and biased signaling. This largely conserved residue^[47]^ restrains the inactive state of several receptors. ^[3,48,49]^ A systematic mutational investigation of the melanocortin 4 receptor indicated its role in signaling bias.^[50]^ We show here that the I130^3.43^N mutation in V2R led to constitutive cAMP signaling without βArr2 recruitment, in line with previous reports.^[31]^

V2R^K100R^, allowing the assignment of K64 by comparison with the spectrum of V2R^K100R^ (Figure S11). K64^1.59^ NMR correlation peak was different according to the ligand bias, confirming the MD predictions. Indeed, the R64^1.59^–Y325^7.53^ Cα distance distribution profile of AVP-V2R retains a component similar to the apo state, contrary to MCF14-V2R. Similarly, the NMR correlation peak of K64^1.59^ in the AVP-bound state is closer to that in the apo state, compared to the MCF14-bound state. (Figure 4).

By contrast, upon AVP stimulation, the V2R^I130N^ variant displayed a higher βArr2 recruitment level than wt V2R (Figure S7), suggesting a particular role of residue I130^3.43^ for βArr2 signaling mediated by its direct interactions with the NP^7.50^xxY motif in TM7. Increasing evidence points to the particular role of TM7 and H8 in the G protein/β-arrestin bias of class A GPCRs. Some studies attributed ligand bias to specific ligand-TM7 interactions,^[51–53]^ others associated biased signaling to distinct conformations in the Na^+^-binding site, the NP^7.50^xxY motif, as well as H8.^[10,11,13,14,18,53– 59]^ In line with previous studies, our work on the V2R reaffirm the presence of a conserved mechanisms associated with biased activation of class A GPCRs (Figure 6). However, the detailed structural features may vary among different receptors, ligands and mutants. Nevertheless, analysis of receptor-ligand interactions and bias-associated receptor conformations can provide direct information for structure-based design of biased ligands.^[18,51,52,55,60]^

## Conclusion

Clinical translation of GPCR ligand bias is challenging due to the complexity of GPCR signaling. The approach of combining ligand paramagnetic tagging with enhanced-sampling MD simulations offers a solution to the common challenge of determining ligand binding poses. Our findings indicate conserved structural patterns for G protein bias that may be common to other class A GPCRs.

## Supporting information

1.Supplemental Experimental Procedures 2.Supplemental Items Table S1. Ligand binding affinity for wt V2R and variants. Table S2. Ligand binding affi

## Supporting Information

The authors have cited additional references within the Supporting Information.

## Authors Contributions

Conceptualization, HD, BM, and XC; investigation, GG, AF, BM, HO, XC, HD, CM, AK, RL, JN, SR, SU, and JD; writing, original draft, XC, BM, SU, DB and HD. writing—review and editing, GG, AF, BM, HO, XC, HD, RS, SG; All authors read and approved the final manuscript.

## Acknowledgements

The CBS belongs to the French Infrastructure for Integrated Structural Biology (FRISBI) and is a GIS-IBISA plaform, supported by the National Research Agency (ANR-10-INBS-05). We also thank the Institut de Génomique Fonctionnelle Arpege Pharmacology platform (http://www.arpege.cnrs.fr) for access to their instruments, and Perkin Elmer CisBio for providing reagents. MS experiments were performed at the Montpellier Proteomics Platform (PPM, BioCampus Montpellier). This work was supported in part by grants from ANR (22-CE44-0021) and FRM (DEQ20150331736). This work was also supported by the University of Strasbourg, the Interdisciplinary Thematic Institute IMS (ANR-10-IDEX-0002) and SFRI (STRAT’US project, ANR-20-SFRI-0012) under the framework of the French Investments for the Future Program. Core funding were provided by CNRS, INSERM and Université de Montpellier. MD simulations were performed using HPC resources from GENCI-TGCC (grant 2021-2022 A0100712461).

## Entry for the Table of Contents

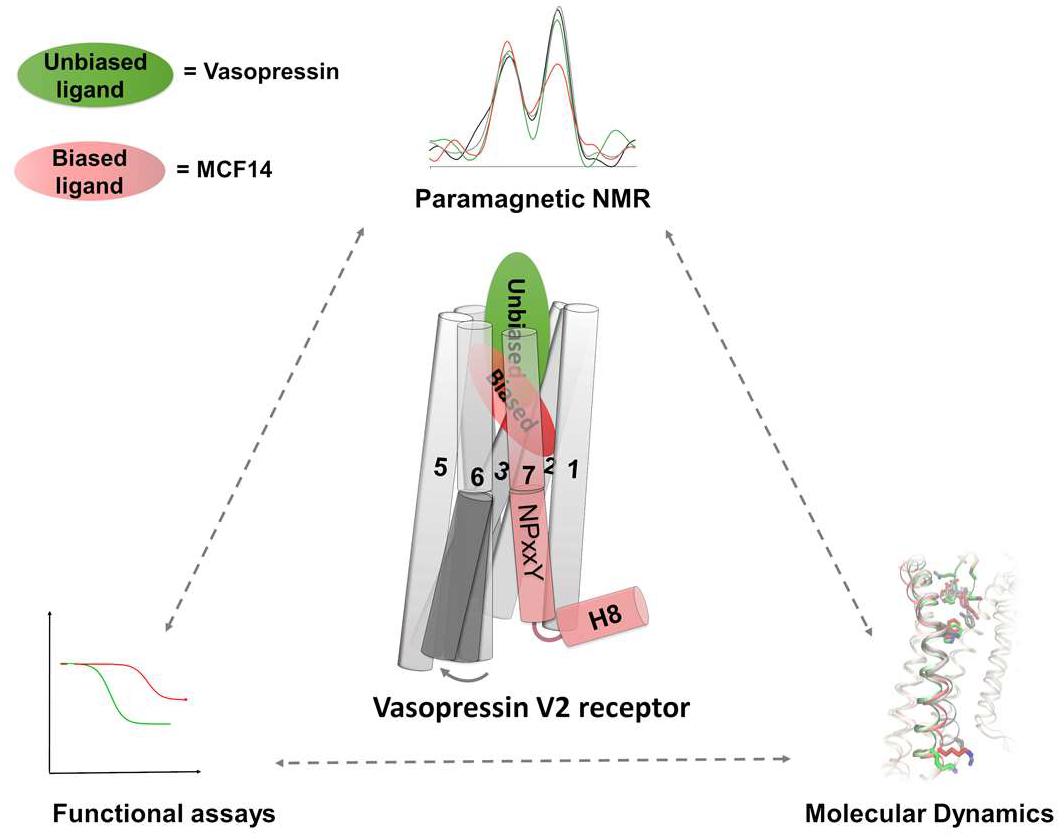

Understanding the structural mechanism of selective biased signalization of G-protein coupled receptors is fundamental in the process of designing safer therapeutic compounds. Taking the vasopressin type 2 receptor as a target, we propose here a unified approach based on a double labelling scheme at the level of the receptor and its ligands using paramagnetic NMR, combined to pharmacological assays and advanced molecular dynamics simulations.

## Notes

Supporting information for this article is given via a link at the end of the document.

### Competing Interest Statement

The authors have declared no competing interest.

## References

[1] D. Wootten, A. Christopoulos, M. Marti-Solano, M. M. Babu, P. M. Sexton, Nat Rev Mol Cell Biol 2018, 19, 638–653.

[2] P. Kolb, T. Kenakin, S. P. H. Alexander, M. Bermudez, L. M. Bohn, C. S. Breinholt, M. Bouvier, S. J. Hill, E. Kostenis, K. A. Martemyanov, R. R. Neubig, H. O. Onaran, S. Rajagopal, B. L. Roth, J. Selent, A. K. Shukla, M. E. Sommer, D. E. Gloriam, Br. J. Pharmacol. 2022, 179, 3651–3674.

[3] Y. X. Tao, Pharmacol. Ther. 2008, 120, 129–148.

[4] L. K. Yang, Z. S. Hou, Y. X. Tao, Biochim. Biophys. Acta - Mol. Basis Dis. 2021, 1867, 165973.

[5] A. J. Venkatakrishnan, X. Deupi, G. Lebon, F. M. Heydenreich, T. Flock, T. Miljus, S. Balaji, M. Bouvier, D. B. Veprintsev, C. G. Tate, G. F. X. Schertler, M. M. Babu, Nat. Publ. Gr. 2016, 536, 484–487.

[6] Q. Zhou, D. Yang, M. Wu, Y. Guo, W. Guo, L. Zhong, X. Cai, A. Dai, W. Jang, E. Shakhnovich, Z. J. Liu, R. C. Stevens, N. A. Lambert, M. M. Babu, M. W. Wang, S. Zhao, Elife 2019, 8, 1–31.

[7] W. I. Weis, B. K. Kobilka, Annu. Rev. Biochem. 2018, 87, 897.

[8] S. G. Rasmussen, H. J. Choi, J. J. Fung, E. Pardon, P. Casarosa, P. S. Chae, B. T. Devree, D. M. Rosenbaum, F. S. Thian, T. S. Kobilka, A. Schnapp, I. Konetzki, R. K. Sunahara, S. H. Gellman, A. Pautsch, J. Steyaert, W. I. Weis, B. K. Kobilka, Nature 2011, 469, 175–180.

[9] S. G. Rasmussen, B. T. DeVree, Y. Zou, A. C. Kruse, K. Y. Chung, T. S. Kobilka, F. S. Thian, P. S. Chae, E. Pardon, D. Calinski, J. M. Mathiesen, S. T. Shah, J. A. Lyons, M. Caffrey, S. H. Gellman, J. Steyaert, G. Skiniotis, W. I. Weis, R. K. Sunahara, B. K. Kobilka, Nature 2011, 477, 549–555.

[10] R. Lamichhane, J. J. Liu, K. L. White, V. Katritch, R. C. Stevens, K. Wüthrich, D. P. Millar, Structure 2020, 28, 371–377.e3.

[11] J. J. Liu, R. Horst, V. Katritch, R. C. Stevens, K. Wuthrich, Science (80-.). 2012, 335, 1106–1110.

[12] M. Louet, M. Casiraghi, M. Damian, M. G. Costa, P. Renault, A. A. Gomes, P. R. Batista, C. M’Kadmi, S. Mary, S. Cantel, S. Denoyelle, K. Ben Haj Salah, D. Perahia, P. M. Bisch, J. A. Fehrentz, L. J. Catoire, N. Floquet, J. L. Banères, Elife 2021, 10, e63201.

[13] R. Rahmeh, M. Damian, M. Cottet, H. Orcel, C. Mendre, T. Durroux, K. S. Sharma, G. Durand, B. Pucci, E. Trinquet, J. M. Zwier, X. Deupi, P. Bron, J.-L. Baneres, B. Mouillac, S. Granier, Proc. Natl. Acad. Sci. 2012, 109, 6733–8.

[14] X. Cong, D. Maurel, H. H. Déméné, I. Vasiliauskaité-brooks, J. Hagelberger, F. Peysson, J. Saint-Paul, J. Golebiowski, S. Granier, R. Sounier, Mol. Cell 2021, 81, 1– 21.

[15] J. Okude, T. Ueda, Y. Kofuku, M. Sato, N. Nobuyama, K. Kondo, Y. Shiraishi, T. Mizumura, K. Onishi, M. Natsume, M. Maeda, H. Tsujishita, T. Kuranaga, M. Inoue, I. Shimada, Angew. Chem. Int. Ed. Engl. 2015, 54, 15771–6.

[16] L. M. Wingler, M. Elgeti, D. Hilger, N. R. Latorraca, M. T. Lerch, D. P. Staus, R. O. Dror, B. K. Kobilka, W. L. Hubbell, R. J. Lefkowitz, Cell 2019, 176, 468–478.e11.

[17] J. Xu, Y. Hu, J. Kaindl, P. Risel, H. Hübner, S. Maeda, X. Niu, H. Li, P. Gmeiner, C. Jin, B. K. Kobilka, Mol. Cell 2019, 75, 53–65.e7.

[18] C. M. Suomivuori, N. R. Latorraca, L. M. Wingler, S. Eismann, M. C. King, A. L. W. Kleinhenz, M. A. Skiba, D. P. Staus, A. C. Kruse, R. J. Lefkowitz, R. O. Dror, Science 2020, 367, 881–887.

[19] Z. Xu, T. Ikuta, K. Kawakami, R. Kise, Y. Qian, R. Xia, M. X. Sun, A. Zhang, C. Guo, X. H. Cai, Z. Huang, A. Inoue, Y. He, Nat. Chem. Biol. 2022, 18, 281–288.

[20] K. Zheng, J. S. Smith, D. S. Eiger, A. Warman, I. Choi, C. C. Honeycutt, N. Boldizsar, J. N. Gundry, T. F. Pack, A. Inoue, M. G. Caron, S. Rajagopal, Sci. Signal. 2022, 15, eabg5203.

[21] J. H. Robben, N. V. A. M. Knoers, P. M. T. Deen, Mol. Biol. Cell 2004, 15, 5693–5699.

[22] X. Ren, E. Reiter, S. Ahn, J. Kim, W. Chen, R. J. Lefkowitz, 2005, 5, 1448–53.

[23] G. Alonso, E. Galibert, V. Boulay, A. Guillou, A. Jean, V. Compan, G. Guillon, Endocrinology 2009, 150, 239–250.

[24] S. G. Ball, Ann. Clin. Biochem. 2007, 44, 417–431.

[25] J. P. Morello, D. G. Bichet, Annu. Rev. Physiol. 2001, 63, 607–630.

[26] D. Bockenhauer, D. G. Bichet, Nat. Rev. Nephrol. 2015, 11, 576–588.

[27] B. Mouillac, C. Mendre, in Target. Traffick. Drug Dev. (Eds.: A. Ulloa-Avuirre, T. Ya-Juong), Springer, 2018, pp. 63–83.

[28] F. Jean-Alphonse, S. Perkovska, M.-C. C. Frantz, T. Durroux, C. Mejean, D. Morin, S. Loison, D. Bonnet, M. Hibert, B. Mouillac, C. Mendre, J Am Soc Nephrol 2009, 20, 2190–2203.

[29] S. S. Rosenthal, B. J. Feldman, G. A. Vargas, S. E. Gitelman, Pediatr Endocrinol Rv 2006, Suppl. 1, 66–70.

[30] J. A. Ballesteros, H. Weinstein, Methods Neurosci. 1995, 25, 366–428.

[31] L. S. Erdélyi, W. A. Mann, D. J. Morris-Rosendahl, U. Groß, M. Nagel, P. Várnai, A. Balla, L. Hunyady, Kidney Int. 2015, 88, 1070–1078.

[32] A. Tiulpakov, C. W. White, R. S. Abhayawardana, H. B. See, A. S. Chan, R. M. Seeber, J. I. Heng, I. Dedov, N. J. Pavlos, K. D. G. Pfleger, Mol. Endocrinol. 2016, 30, 889– 904.

[33] V. Vezzi, C. Ambrosio, M. C. Grò, P. Molinari, G. Süral, T. Costa, H. O. Onaran, S. Cotecchia, Sci. Rep. 2020, 10, 9111.

[34] J. Bous, H. Orcel, N. Floquet, C. Leyrat, J. Lai-Kee-Him, G. Gaibelet, A. Ancelin, J. Saint-Paul, S. Trapani, M. Louet, R. Sounier, H. Déméné, S. Granier, P. Bron, B. Mouillac, Sci. Adv. 2021, 21, eabg5628.

[35] F. Zhou, C. Ye, X. Ma, W. Yin, T. I. Croll, Q. Zhou, X. He, X. Zhang, D. Yang, P. Wang, H. E. Xu, M. W. Wang, Y. Jiang, Cell Res. 2021, 31, 929–931.

[36] L. Wang, J. Xu, S. Cao, D. Sun, H. Liu, Q. Lu, Z. Liu, Y. Du, C. Zhang, Cell Res. 2021, 31, 932–934.

[37] J. Bous, A. Fouillen, H. Orcel, S. Trapani, X. Cong, S. Fontanel, J. Saint-Paul, J. Lai-Kee-Him, S. Urbach, N. Sibille, R. Sounier, S. Granier, B. Mouillac, P. Bron, Sci. Adv. 2022, 8, eabo7761.

[38] M. P. Bokoch, Y. Zou, S. G. F. Rasmussen, C. W. Liu, R. Nygaard, D. M. Rosenbaum, J. J. Fung, H. J. Choi, F. S. Thian, T. S. Kobilka, J. D. Puglisi, W. I. Weis, L. Pardo, R. S. Prosser, L. Mueller, B. K. Kobilka, Nature 2010, 463, 108–112.

[39] R. Sounier, C. Mas, J. Steyaert, T. Laeremans, A. Manglik, W. Huang, B. K. Kobilka, H. Déméné, S. Granier, Nature 2015, 524, 375–378.

[40] R. D. J. Kimbrough, W. D. Cash, L. A. Branda, W. Chan, V. Du Vigneau, J. Biol. Chem. 1963, 238, 1411–4.

[41] H. Tabata, T. Yoneda, T. Oshitari, H. Takahashi, H. Natsugari, J. Med. Chem. 2017, 60, 4503–4509.

[42] R. H. Oakley, S. A. Laporte, J. A. Holt, L. S. Barak, M. G. Caron, J. Biol. Chem. 1999, 274, 32248–32257.

[43] J. Orloff, J. S. Handler, J. Clin. Invest. 1962, 41, 702–709.

[44] Y. Yamamura, S. Nakamura, S. Itoh, T. Hirano, T. Onogawa, T. Yamashita, Y. Yamada, K. Tsujimae, M. Aoyama, K. Kotosai, H. Ogawa, H. Yamashita, K. Kondo, M. Tominaga, G. Tsujimoto, T. Mori, J Pharmacol Exp Ther 1998, 287, 860–867.

[45] S. K. Huang, A. Pandey, D. P. Tran, N. L. Villanueva, A. Kitao, R. K. Sunahara, A. Sljoka, R. S. Prosser, Cell 2021, 184, 1884–1894.e14.

[46] A. El Daibani, J. M. Paggi, K. Kim, Y. D. Laloudakis, P. Popov, S. M. Bernhard, B. E. Krumm, R. H. J. Olsen, J. Diberto, F. I. Carroll, V. Katritch, B. Wünsch, R. O. Dror, T. Che, Nat. Commun. 2023, 14, 1338.

[47] T. Mirzadegan, G. Benko, S. Filipek, K. Palczewski, Biochemistry 2003, 42, 2759.

[48] S. Kosugi, N. Hai, H. Okamoto, H. Sugawa, T. Mori, Eur. J. Endocrinol. 2000, 143, 471–477.

[49] E. Ceraudo, M. Horioka, J. M. Mattheisen, T. D. Hitchman, A. R. Moore, M. A. Kazmi, P. Chi, Y. Chen, T. P. Sakmar, T. Huber, J. Biol. Chem. 2021, 296, 100163.

[50] X. L. Mo, R. Yang, Y. X. Tao, J. Mol. Endocrinol. 2012, 49, 221–235.

[51] J. D. McCorvy, D. Wacker, S. Wang, B. Agegnehu, J. Liu, K. Lansu, A. R. Tribo, R. H. J. Olsen, T. Che, J. Jin, B. L. Roth, Nat Struct Mol Biol 2018, 25, 787–796.

[52] D. Cao, J. Yu, H. Wang, Z. Luo, X. Liu, L. He, J. Qi, L. Fan, L. Tang, Z. Chen, J. Li, J. Cheng, S. Wang, Science 2022, 375, 403–411.

[53] Z. Shao, Q. Shen, B. Yao, C. Mao, L. N. Chen, H. Zhang, D. D. Shen, C. Zhang, W. Li, X. Du, F. Li, H. Ma, Z. H. Chen, H. E. Xu, S. Ying, Y. Zhang, H. Shen, Nat. Chem. Biol. 2022, 18, 264–71.

[54] D. Wacker, C. Wang, V. Katritch, G. W. Han, X. P. Huang, E. Vardy, J. D. McCorvy, Y. Jiang, M. Chu, F. Y. Siu, W. Liu, H. E. Xu, V. Cherezov, B. L. Roth, R. C. Stevens, Science (80-.). 2013, 340, 615–619.

[55] J. D. McCorvy, K. V. Butler, B. Kelly, K. Rechsteiner, J. Karpiak, R. M. Betz, B. L. Kormos, B. K. Shoichet, R. O. Dror, J. Jin, B. L. Roth, Nat. Chem. Biol. 2018, 14, 126– 134.

[56] P. Xu, S. Huang, H. Zhang, C. Mao, X. E. Zhou, X. Cheng, I. A. Simon, D. D. Shen, H. Y. Yen, C. V. Robinson, K. Harpsøe, B. Svensson, J. Guo, H. Jiang, D. E. Gloriam, K. Melcher, Y. Jiang, Y. Zhang, H. E. Xu, Nature 2021, 592, 469–473.

[57] Y. Peng, J. D. McCorvy, K. Harpsøe, K. Lansu, S. Yuan, P. Popov, L. Qu, M. Pu, T. Che, L. F. Nikolajsen, X. P. Huang, Y. Wu, L. Shen, W. E. Bjørn-Yoshimoto, K. Ding, D. Wacker, G. W. Han, J. Cheng, V. Katritch, A. A. Jensen, M. A. Hanson, S. Zhao, D. E. Gloriam, B. L. Roth, R. C. Stevens, Z. J. Liu, Cell 2018, 172, 719–730.e14.

[58] A. M. Schönegge, J. Gallion, L. P. Picard, A. D. Wilkins, C. Le Gouill, M. Audet, W. Stallaert, M. J. Lohse, M. Kimmel, O. Lichtarge, M. Bouvier, Nat. Commun. 2017, 8, 2169.

[59] L. M. Wingler, C. McMahon, D. P. Staus, R. J. Lefkowitz, A. C. Kruse, Cell 2019, 1–12.

[60] H. Wang, F. Hetzer, W. Huang, Q. Qu, J. Meyerowitz, J. Kaindl, H. Hübner, G. Skiniotis, B. K. Kobilka, P. Gmeiner, Angew. Chem. Int. Ed. Engl. 2022, 61, e202200269.

