## Supplementary material for "Biased activation of the vasopressin V2 receptor probed by NMR, paramagnetic ligands, and molecular dynamics simulations": 1.Supplemental Experimental Procedures 2.Supplemental Items Table S1. Ligand binding affinity for wt V2R and variants. Table S2. Ligand binding affi

#### Table of Contents

1. **Supplemental Experimental Procedures**
2. **Supplemental Items**

Table S1. Ligand binding affinity for wt V2R and variants.
Table S2. Ligand binding affinity and potency for V2R$^{K100R}$ variant.
Scheme S1. Synthesis of paramagnetic ligands.
Figure S1. Engineering of V2R and ligands.
Figure S2. Analysis of lysine methylation by NMR and mass spectrometry**.**
Figure S3. Assignment of correlation peaks of dimethylated lysines K100, K116 and K268.
Figure S4. Effect of $^{par}$MCF14 binding and reduction by ascorbic acid on the lysine HMQC correlation peaks of wt V2R and fitting procedure.
Figure S5. Conformational dynamics of V2R and ligands during MD simulations.
Figure S6. Distances between the paramagnetic tag and dimethylated K100$^{2.65}$ and K116$^{3.29}$ during the MD simulations.
Figure S7. Pharmacological and functional properties of the mutant V2R-I130N.
Figure S8. Probability density distribution of the conformational features measured during the MD simulations.
Figure S9. Probability density map of two conformational features during the MD simulations.
Figure S10. Functional properties and NMR spectra of the V2R$^{K100R}$ mutant
Figure S11. Assignment of correlation peaks of the probe K64 of V2$^{R64K-K100R}$.
Figure S12. Scheme of REST2 protocol.

3. **Supplemental References**

### 1. Supplemental Experimental Procedures

#### V2R construction and expression.

The V2R construct used for NMR analysis was already described.[1] It is also shown in Figure S1A. Briefly, the *Spodoptera frugiperda (Sf9)*-optimized sequence of the human V2R was cloned into a pFastBac™1 vector (Invitrogen) using EcoR1/Xba1 restriction sites. To facilitate expression and purification of the V2R construct, the hemagglutinin signal peptide (MKTIIALSYIFCLVFA) followed by a Flag tag (DYKDDDDA) were added at the N-terminus, and a Twin-strep-tag® (WSHPQFEKGGGSGGGSGGGSWSHPQFEK) inserted at the C-terminus. In addition, N22 was substituted with a glutamine residue to avoid N-glycosylation, and C358 mutated into an alanine to eliminate potential intermolecular disulfide bridges during solubilization and purification. A Tobacco Etch Virus (TEV) protease cleavage site (ENLYFQG - following the Flag tag) and 2 Human Rhinovirus 3C protease (3C) cleavage sites (LEVLFQGP - 1 inserted in the N-terminus between D30 and T31, the other inserted in the C-terminus between G345 and Q354 and replacing R346-TPPSLG-P353) were also added to remove N and C termini and facilitate NMR analysis (Figure S1A). After cleavage, the resulting sequence corresponds to a 323 amino-acid residue protein with a theoretical 35.5 kDa molecular weight. M1L2 residues were replaced by AS residues, and LE residues were added before the Twin-strep-tag®, during subcloning (introduction of Nhe1 and Xho1 restriction sites, respectively). Sequence modifications did not affect the receptor ligand binding or function.[1] The V2R was expressed in Sf9 insect cells using the Bac-to-Bac® baculovirus expression system (Invitrogen) according to manufacturer's instructions. Insect cells were grown in suspension in EX-CELL® 420 medium (Sigma-Aldrich) to a density of 4 × $10^6$ cells per ml and infected with the recombinant baculovirus encoding V2R at a multiplicity of infection of 2 to 3. The culture medium was supplemented with the specific V2R pharmacochaperone antagonist Tolvaptan (Sigma-Aldrich) at 1 µM to allow proper folding and membrane targeting of the receptor.[2,3] The cells were infected for 48 h to 54 h at 28 °C. Before harvesting, the expression of the V2R was controlled by immunofluorescence using an anti-Flag M1 antibody coupled to an Alexa488 fluorophore. Cells were then harvested by centrifugation (2 steps, 15 min then 20 min at 3,000g) and the cell pellets were stored at -80 °C until use.

The different mutants of the V2R ($V2R^{K100R}$, $V2R^{K116R}$, $V2R^{K268R}$, $V2^{R64K\text{-}K100R}$) were all derived from the matrix described above using the gene synthesis and molecular biology services of Eurofins Genomics.

#### V2R purification and labeling.

After thawing the frozen cell pellets, cells were lysed by osmotic shock in 10 mM Tris-HCl pH 8, 1 mM EDTA buffer containing 2 $mg.ml^{-1}$ iodoacetamide (Sigma-Aldrich), Tolvaptan 1 µM and protease inhibitors [leupeptine (5 $µg.ml^{-1}$) (Euromedex), benzamidine (10 $µg.ml^{-1}$) (Sigma-Aldrich), and phenylmethylsulfonyl fluoride (PMSF) (10 $µg.ml^{-1}$) (Euromedex)]. Lysed cells were centrifuged (15 min at 38,400g) and the pellets containing crude membranes were solubilized using a glass dounce tissue grinder (15 + 20 strokes using A and B pestles respectively) in a solubilization buffer containing 20 mM Tris-HCl pH 8, 500 mM NaCl, 0.5 % (w/v) n-dodecyl-β-D-maltopyranoside (DDM, Anatrace), 0.2 % (w/v) sodium cholate (Sigma-Aldrich), 0.03 % (w/v) cholesteryl hemisuccinate (CHS, Sigma-Aldrich), 20 % Glycerol (VWR), 2 $mg.ml^{-1}$ iodoacetamide, 0.75 $ml.L^{-1}$ Biotin Biolock (IBA), Tolvaptan 1 µM and protease inhibitors. The extraction mixture was stirred for 1 h at 4 °C and centrifuged (20 min at 38,400g).

The cleared supernatant was poured onto equilibrated Strep-Tactin resin (IBA) for a first affinity purification step. After 2 h of incubation at 4 °C under stirring, the resin was laid down into a column and washed three times with 10 column volume (CV) of a buffer containing 20 mM Tris-HCl pH 8, 500 mM NaCl, 0.1 % (w/v) DDM, 0.02 % (w/v) sodium cholate, 0.03 % (w/v) CHS, Tolvaptan 1 µM. The bound receptor was eluted in the same buffer supplemented with 2.5 mM Desthiobiotin (IBA). The amount of V2R was calculated by UV absorbance spectroscopy.

The eluate supplemented with 2 mM $CaCl_2$ was loaded onto an M1 anti-Flag affinity resin (Sigma-Aldrich) for a second affinity purification. After loading, the DDM detergent was then gradually exchanged into a buffer with Lauryl Maltose Neopentyl glycol (LMNG, Anatrace) (20 mM HEPES pH 7.4, 100 mM NaCl, 0.01 % CHS and 0.5 % LMNG). The LMNG concentration was then decreased from 0.5 % to 0.02 %. The V2R was then eluted in 20 mM HEPES pH 7.4, 100 mM NaCl, 0.02 % LMNG, 0.002 % CHS, 2 mM EDTA, 10 µM Tolvaptan and 0.2 $mg.ml^{-1}$ Flag peptide (Covalab). The absence of tolvaptan during the SEC, the two dialysis steps and during the final dilution /concentration of the sample ensured that the orthosteric binding site of the V2R was empty and available for measuring effects of the different biased and unbiased ligands. The amount of V2R was estimated by UV absorbance spectroscopy and was then cleaved overnight using the HRV3C protease at a 1:20 weight ratio (HRV3C:V2R) at 4 °C. Concomitantly, the V2R was labeled onto its lysine residues by reductive methylation using 10 mM $^{13}C$-formaldehyde (Sigma-Aldrich) in the presence of 10 mM sodium cyanoborohydride (Sigma-Aldrich) following the protocol described previously.[4,5]

The 3C-cleaved $^{13}C$-dimethyl-lysine-labeled V2R was concentrated using 50-kDa concentrators (Millipore) and separated from the protease using a Superdex 200 10/300 column on an AKTA purifier system (0.5 mL/min flowrate, buffer 20 mM HEPES pH 7.4, 100 mM NaCl, 0.02 % MNG, no ligand). Fractions corresponding to the pure monomeric V2R were pooled and dialyzed first 2 h at 4 °C using a 3-kDa MWCO cassette in 20 mM HEPES pH 7.4, 40 mM NaCl, 0.02 % LMNG, 0.002 % CHS. Then, the sample was dialyzed again 2 h at 4 °C in 98.85 % $D_2O$ buffer with 20 mM HEPES-d18 pH 7.4 (uncorrected), 40 mM NaCl, 0.02 % LMNG, 0.002 % CHS.

V2R was then concentrated using a 50-kDa MWCO concentrator up to 250 µl, and then diluted with 2 ml of 98.85 % $D_2O$ buffer with 20 mM HEPES-d18 pH 7.4, 40 mM NaCl before a second concentration step. The final concentration of V2R was 25-30 µM in buffer containing 20 mM HEPES-d18 pH 7.4, 40 mM NaCl, 0.0022 % LMNG and 0.0002 % CHS (NMR buffer). Acid 4,4-dimethyl-4-silapentane-1-sulfonique (DSS) was added at a final concentration of 100 µM as a standard to calibrate the NMR analysis.

#### Mass spectrometry and NMR analysis of lysine methylation.

Samples were prepared as previously described.[6] We used $V2R^{K100R}$ to better assign the NMR signals from $K268^{6.32}$. Briefly, to perform these experiments, the $V2R^{K100R}$ mutant was expressed, extracted and purified. Following the M1 anti-flag affinity chromatography, the receptor was cleaved by the HRV3C protease and concomitantly labelled (or not) onto its lysine residues by reductive methylation. At this step, only two residues can be labelled, K116 and K268 (see Figure S1A). Then, the samples were concentrated, subjected to size exclusion chromatography, and the eluted fractions concentrated to 1 µg/µl. The purified $V2R^{K100R}$, labelled or unlabelled (10 µg per sample), was digested using micro S-Trap columns (https://protifi.com/; Huntington, NY) following the supplier's protocol using 1 µg of trypsin (Promega, Gold) for 2 hours at 47°C. The peptides obtained were analyzed using nano-throughput high-performance liquid chromatography (Ultimate 3000-RSLC, Thermo Fisher Scientific) coupled to a mass spectrometer (Q Exactive-HF, Thermo Fisher Scientific) equipped with a nanospray source. Raw spectra were processed using the MaxQuant v2.0.3.0.[7] Signal intensities of receptor peptides were extracted using Skyline v21.1.0.245.[8] Graphical representation of the spectra was performed using FreeStyle v1.8 SP2 (Thermo Fisher Scientific Inc.). The comparison of peptides from unlabeled and labeled $V2R^{K100R}$ allowed to conclude that reductive dimethylation is nearly complete with more than 95% efficiency (Figure S2B-C). Consistent with these results, no monomethylated peak could be in found in the 2,4-3 ppm/34.5-36.5 ppm ($^1H/^{13}C$) region of HMQC spectra of native and mutant receptors (Figure S2A).

#### NMR spectroscopy.

Final samples (~270 µl at 30-35µM) were loaded into Shigemi microtubes whose susceptibility matched with $D_2O$. All data for ligands and mutant studies were acquired on 700 MHz Bruker Avance III spectrometers (Bruker, Rheinstetten, Germany), equipped with 5 mm cryogenic H/C/N/D probes with z axis gradient at 293K, unless otherwise specified. $^1H$-$^{13}C$ correlation spectra were recorded using heteronuclear multiple-quantum coherence (HMQC) experiments in echo anti-echo mode. Spectral widths in ω1 and ω2 were 8,417.5 Hz and 3,518.6 Hz at 700 MHz centred at 40 p.p.m. $^{13}C$ decoupling was performed with a GARP4 sequence. Thirty-two steady-state scans preceded data acquisition. Typically, 134 complex points with per FID were recorded, to ensure a 27-Hz resolution per point at 700 MHz before zero filling, with 32 scans and a relaxation delay of 1.5 s ($^{par}$dLVP, $^{par}$MCF14) or 80 scans and a relaxation delay of 0.5 s (AVP, MCF14, TVP). Total collection time varied between 3 and 4 h, depending on the sample concentration. Ligands were dissolved in NMR buffer (AVP, $^{par}$dLVP) or in perdeuterated dimethyl d6-sulfoxide (d6-DMSO, Cambridge Isotope) (MCF14, $^{par}$MCF14, TVP) to 10 mM and directly added to the sample in the Shigemi tube at a final concentration five (AVP, MCF14) or 0.95 fold ($^{par}$dLVP, $^{par}$MCF14) of receptor. The non-paramagnetic agonists (AVP, MCF14, TVP) have high affinities for V2R (< 0.1 µM). At saturating concentrations (30 µM and 150 µM for V2R and non-paramagnetic ligands, respectively), according to the law of mass action, 99% of the V2R should be in complex at 10 µM concentration of agonists (~100 times the Kd concentration). The effect of paramagnetic ligands was studied using sub-stoichiometric ligand concentrations, to promote binding in the receptor orthosteric site. Reduction was achieved by the addition of 5 equivalents of freshly prepared ascorbic acid dissolved at 10 mM in the NMR buffer.

All NMR spectra were processed using the suite of programs provided in the NMRPipe/NMRDraw software distribution.[9] The spectra were normalized using DSS (2,2-dimethyl-2-silapentane-5-sulfonic acid) as an internal reference. For peak fitting analysis with the NMRPipe sofware, spectra were processed with a Gaussian window, and zero-filled to 4096 x 1024 data points in time domain data t2 and t1, respectively. Briefly, the g1 parameter corresponding to line sharpening was adjusted so that the time-domain signal no longer decays. The g2 parameter corresponding to line broadening was then selected, with the goal of getting an exponentially well-decayed FID after both g1 and g2 are applied together. Peaks of processed spectra were then fitted with the program nlinLS, provided as a part of the NMRPipe packagestarting from values obtained from the peak-peaking routine. The nlinLS explicitely takes into account the overlap of peaks, which are fitted simultaneously and not independently by adjusting the parameter Clustid. The quality of the fits was examined visually by estimating the residual difference between the experimental data and the results of the model calculations (Figure S4B). Peak intensity of the $^{13}C$-dimethylated lysines were normalized to the intensity of the G52 (cleaved N terminus) and buffer peaks. Errors in the peak volume were calculated replicating experiments starting purification of new V2R culture batches (AVP, n=3; $^{par}$AVP, n=2; MCF14, n=4; $^{par}$MCF14, n=2; TVP, n=3). Figures were drawn with the Topspin 3.6 (Bruker, Inc) and NMRview packages.[10]

#### Time-resolved Fluorescence Resonance Energy Transfer binding assays.

V2R binding studies using Tag-Lite assays (PerkinElmer Cisbio) based on time-resolved fluorescence resonance energy transfer (TR-FRET) measurements were previously described.[1,11,12] The technology combines the advantages of FRET with time-resolved measurement of fluorescence, eliminating short-lived background fluorescence. In binding studies, the V2R is labeled with the donor acceptor, an europium cryptate (Eu cryptate), and the ligand is labeled with a fluorescence red acceptor, the d2 (DY647). The donor emits fluorescence upon excitation and the energy is transferred to the acceptor if binding occurs, leading to a specific fluorescence signal. The donor fluorescence is also measured and a ratio with the fluorescence applied (665 nm/620 nm, see below). Briefly, HEK cells were plated (15,000 per well) in precoated with poly-L-ornithine (14 μg/ml, Sigma-Aldrich) white-walled, flat-bottom, 96-well plates (Greiner CELLSTAR plate, Sigma-Aldrich) in Dulbecco's minimum essential medium (DMEM) containing 10 % fetal bovine serum (FBS, Eurobio), 1 % nonessential amino acids (GIBCO), and penicillin/streptomycin (GIBCO). Cells were transfected 24 h later with a plasmid coding for the 3C-cleaved V2R version used in NMR studies (Figure S1A) fused at its N-terminus to the enzyme-based self-labeling SNAP-tag (pRK5-SNAP vector, PerkinElmer Cisbio). The I130N mutation was introduced in this V2R construction (Figure S1A). Transfections were performed with X-tremeGENE 360 (Merck), according to the manufacturer's recommendations: 10 μl of a premix containing DMEM, X-tremeGENE 360 (0.3 μl per well), SNAP-V2 coding plasmid (30 ng per well, whatever the construct used), and noncoding plasmid (70 ng per well) were added to the culture medium. After a 48-hour culture period, cells were rinsed once with Tag-lite medium (PerkinElmer Cisbio) and incubated in the presence of Tag-lite medium containing 100 nM benzylguanine-Lumi4-Tb for at least 60 min at 37 °C. Cells were then washed four times. For saturation studies, cells were incubated for at least 4 h at 4 °C in the presence of benzazepine-red nonpeptide vasopressin antagonist (BZ-DY647, PerkinElmer Cisbio) at various concentrations ranging from $10^{-10}$ to $10^{-7}$ M. Nonspecific binding was determined in the presence of 10 μM vasopressin. For competition studies, cells were incubated for at least 4 h at 4 °C with benzazepine-red ligand (5 nM) and increasing concentrations of vasopressin ranging from $10^{-12}$ to $10^{-5}$ M. Fluorescent signals were measured at 620 nm (fluorescence of the donor) and at 665 nM (FRET signal) on a PHERAstar (BMG LABTECH). Results were expressed as the 665/620 ratio [10,000 × (665/620)]. A specific variation of the FRET ratio was plotted as a function of benzazepine-red concentration (saturation experiments) or competitor concentration (competition experiment). All binding data were analyzed with GraphPad 9.1.1 (GraphPad Prism Software Inc.) using the one site-specific binding equation. All results are expressed as the means ± SEM of at least three independent experiments performed in triplicate. $K_i$ values were calculated from median inhibitory concentration values with the Cheng-Prusoff equation.

#### cAMP accumulation assays.

V2R (wt and mutants $V2R^{K100R}$, $V2R^{I130N}$) functional studies based on TR-FRET measurements were described previously.[1,2,13] Briefly, HEK cells were plated in pre-coated black-walled 96-well plates (Falcon) at 5,000 cells per well, and then transfected 24 h later with a plasmid coding for the SNAP-tagged V2R version used in NMR studies. Transfections were performed with X-tremeGENE 360 (Merck), according to the manufacturer's recommendations: 10 μl of a premix containing DMEM, X-tremeGENE 360 (0.3 μl per well), pRK5-SNAP-V2R coding plasmid (ranging from 0.3 to 30 ng per well, depending on the V2R construct), and pRK5 noncoding plasmid (ranging from 70 to 99.7 ng per well) were added to the culture medium. After a 24-hour culture period, cells were treated for 30 min at 37 °C in the cAMP buffer with or without increasing ligands concentrations ($10^{-12}$ to $3.16 \times 10^{-5}$ M) in the presence of 0.1 mM RO201724, a phosphodiesterase inhibitor (Sigma-Aldrich). The accumulated cAMP was quantified using the cAMP Dynamic 2 Kit (PerkinElmer Cisbio) according to the manufacturer's protocol. Native cAMP produced by cells upon activation competes with d2-labeled cAMP (red acceptor) for binding to monoclonal Eu cryptate-labeled antibody (fluorescent donor). The specific signal is inversely proportional to the concentration of cAMP (in the dose-response experiments, the FRET signal is thus decreasing as a function of cAMP increase in the sample). Fluorescent signals were measured at 620 and 665 nm on a PHERAstar microplate reader. Data were plotted as the FRET ratio [10,000 × (665/620)] as a function of AVP concentration [log (AVP)]. Data were analyzed with GraphPad Prism v 9.1.1 (GraphPad Prism Software Inc.) using the "dose-response stimulation" subroutine. Median effective concentrations were determined using the log(agonist) versus response variable slope (four parameters) fit procedure. Experiments were repeated at least three times on different cultures, each condition in triplicate. Data are presented as means ± SEM.

#### β-arrestin2 recruitment assays.

The recruitment assay of βArr2 was previously described.[6] Briefly, upon GPCR activation, βArrs are recruited to stop G protein signaling and to initiate clathrin-mediated receptor internalization. During this process, the release of the C-terminal domain of βArrs is associated with the binding of βArrs to the adaptor protein 2 (AP2). This interaction can be measured using the HTRF® technology (PerkinElmer CisBio) based on the use of two specific antibodies, one directed against βArr2, the second one specific for AP2. In this assay (βArr2 recruitment kit, PerkinElmer CisBio), the AP2 antibody is labeled with an Eu cryptate fluorescent donor, and the one against βArr2 is labeled with a d2 fluorescent acceptor, their proximity being detected by FRET signals. The specific signal is positively modulated in proportion with the recruitment of βArr2 to AP2 upon V2R activation by AVP. Briefly, HEK cells were plated at a seeding density of 25,000 cells per well in a precoated white-walled 96-well plates (CELLSTAR plate, Sigma-Aldrich) for 24 h, in DMEM complemented with 10 % FBS, 1 % non-essential amino acids, and 1 % penicillin-streptomycin antibiotics solution. To produce the V2R, the cells were transfected with a pRK5-SNAP-V2R plasmid (from 0.6 to 30 ng per well, depending on the construct, wt or I130N NMR versions) using X-tremeGENE 360 (Merck), according to the manufacturer's recommendations. After a 24-hour culture, cells were used to evaluate the recruitment of βArr2 to AP2 upon V2R activation with the βArr2 recruitment kit (PerkinElmer CisBio) following the manufacturer recommendation. Briefly, the cells were first washed one time with DMEM-free and incubated 2 h at room temperature (RT) with 100 μl per well of stimulation buffer containing various concentrations of the different ligands (ranging from $10^{-12}$ M to $10^{-5}$ M).

The medium was then replaced by 30 µl per well of stabilization buffer for 15 min at RT. The cells were then washed three times with 100 µl per well of wash buffer before adding 100 µl per well of a pre-mix of Eu cryptate and d2 antibodies in detection buffer. Following overnight incubation at RT, 80 µl of medium was removed from each well before reading the 96-well plates a PHERAstar by measuring the signals of the donor (Eu cryptate-labeled AP2 antibody) at a wavelength of 620 nm, and of the acceptor at 665 nm (d2-labeled βArr2). Finally, the results were expressed as the FRET ratio [(665/620) x 10,000] and plotted using GraphPad 9.1.1 (GraphPad Prism software inc.). Experiments were repeated at least three times on different cultures, each condition in triplicate. Data are presented as means ± SEM.

#### Statistical analysis

As indicated in TR-FRET binding assays, cAMP accumulation assays and in β-arrestin2 recruitment assays, values and SEM were calculated form at least 3 independent experiments. The p values were obtained by T-test analysis of each condition from data in Excel (Microsoft Windows, Albuquerque, NM, USA). Statistical significance was defined as ns, $p > 0.05$; *$p < 0.05$; **$p < 0.01$; ***$p < 0.001$; and ****$p < 0.0001$.

#### ~~Supplementary methods for~~ Ligand chemistry

**General**

Reagents were obtained from commercial sources and used without any further purification. (Deamino-Cys$^1$, Lys$^8$)-Vasopressin trifluoroacetate was purchased from Bachem. Compound 1 was synthesized as previously described.[14] Thin-layer chromatography was performed on Merck silica gel 60F254 plates. VWR silica gel (40-63 µm) was used for chromatography columns. Semi-preparative reverse-phase HPLC purifications were performed on a Waters SunFire C18 OBD Prep column (5 µm, 19 × 150 mm) on a Gilson PLC2020 system. Analytical reverse-phase HPLC were performed on a Ascentis C18 column (2.7 µm, 7.5 cm × 4.6 mm) on an Agilent Technologies 1200 series HPLC system using a linear gradient (5% to 100% v/v in 7.3 min, flow rate of 1.6 mL.min-1) of solvent B (0.1% v/v TFA in $CH_3CN$) in solvent A (0.1% v/v TFA in $H_2O$). $^1$H NMR spectra were recorded at 400 MHz, $^{13}$C NMR spectra were recorded at 101 MHz on a Bruker Advance spectrometer. Chemical shifts are reported in parts per million (ppm), and coupling constants (J) are reported in hertz (Hz). Signals are described as s (singlet), d (doublet), t (triplet), q (quadruplet), p (pentuplet) and m (multiplet). High-resolution mass spectra (HRMS) were obtained on an Agilent Technologie 6520 Accurare-Mass Q.Tof LC/MS apparatus equipped with a Zorbax SB C18 column (1.8 µm, 2.1 × 50 mm) using electrospray ionization (ESI) and a time-of-flight analyzer (TOF).

**Synthesis of paramagnetic ligands**

*$^{par}$MCF14*

The synthesis of $^{par}$MCF14 was performed in two steps (see Scheme S1A).

Tert-butyl (6-((1-(2-chloro-4-(3-methyl-1H-pyrazol-1-yl) benzoyl)-2,3,4,5-tetrahydro-1H-benzo[b]azepin-5-yl) oxy)hexyl)carbamate (compound 2). To a solution of 1 (281.6 mg, 0.74 mmol) in 20 mL of anhydrous DMF at 0°C under argon atmosphere was added NaH 60% (70.7 mg, 1.84 mmol) portion wise. After 15 min, tert-butyl N-(6-bromohexyl) carbamate (517 mg, 1.84 mmol) in 8 mL of anhydrous DMF was added dropwise and stirred for 18 h at room temperature. The crude product was diluted in saturated aqueous $NaHCO_3$ and extracted with $CH_2Cl_2$. The organic phases were dried over $Na_2SO_4$, and evaporated. The residue was purified by chromatography on a silica gel column (0% to 30% EtOAc in *n*-heptane) to afford a clear oil (307 mg; yield 72%). $t_R$ = 6.71 min. $^1$H RMN (400 MHz, DMSO-*d$_6$*) : *δ* 8.39-8.37 (m, 1H), 8.04-7.71 (m, 2H), 7.41-7.25 (m, 3 H), 7.11-6.96 (m, 2H), 6.83-6.71(m, 1H), 6.32-6.31 (m, 1H), 4.79-4.74 (m, 1H), 4.53-4.52 (m, 1H), 3.57-3.38 (m, 2H), 2.94-2.73 (m, 3H), 2.37-2.14 (m, 4H), 2.02-1.98 (m, 1H), 1.69-1.52 (m, 4H), 1.41-1.20 (m, 17H). $^{13}$C (100 MHz, DMSO-*d$_6$*) : *δ* 165.3, 164.8, 155.5, 150.5, 141.6, 140.1, 139.8, 138.7, 137.4, 133.3, 133.1, 131.5, 130.8, 129.3, 128.8, 128.7, 128.5, 127.7, 127.3, 127.2, 127.1, 117.8, 117.6, 116.5, 115.5, 114.8, 108.5, 81.0, 79.1, 77.7, 77.2, 68.9, 68.2, 46.9, 46.1, 39.8, 32.7, 29.4, 28.2, 26.2, 25.5, 24.9, 22.5, 13.3.

*$^{par}$MCF14.* Compound 2 (14 mg, 0.03 mmol) was dissolved in a solution HCl 4N/Dioxane (400 µL) and was stirred 30 min at room temperature and then evaporated. The residue was dissolved in DMF (1 mL) under argon. Then 3-carboxy-PROXYL (6.7 mg, 0.036 mmol) and PyBOP (18.7 mg, 0.04 mmol) were added to the solution followed by DIEA (0.03 mL, 0.18 mmol). Then the resulting mixture was stirred for 1 h and evaporated. The expected compound was isolated by semi-preparative RP-HPLC on a Sunfire RP-C18 column using a linear gradient of solvent B in solvent A. Fractions containing the product were freeze-dried to afford a clear oil (10.7 mg; yield 55 %). $t_R$ = 5.80 min. HRMS (EI) *m/z* calcd for $C_{36}H_{48}ClN_5O_4$ $[M+H]^+$ 649.3317, found : 649.3305.

*$^{par}$dLVP*

The synthesis of $^{par}$dLVP was performed in one step (see Scheme S1B).

To a solution of (deamino-Cys1, Lys8)-Vasopressin trifluoroacetate (2.5 mg, 2.16 µmol) and 3-carboxy-PROXYL (0.48 mg, 2.6 µmol) in anhydrous DMF (100 µL) under argon atmosphere were added DIEA (2.15 µL, 12.9 µmol) and PyBOP (1.35 mg, 2.56 µmol), stirred for 1 h at room temperature and then evaporated. The expected compound was isolated by semi-preparative RP-HPLC on a Sunfire RP-C18 column using a linear gradient of solvent B in solvent A. Fractions containing the product were lyophilized to afford a white solid (1.7 mg; yield 65%). $t_R$ = 3.15 min. HRMS (EI) *m/z* calcd for $C_{55}H_{79}N_{13}O_{14}S_2$ $[M+H]^+$, 1209.5310, found : 1209.5315.

#### Molecular Dynamic simulations.

*Principle*

REST2[15] is a type of Hamiltonian replica exchange simulation scheme, which performs many replicas of the same MD simulation system in parallel with the original system. The replicas have modified free energy surfaces to facilitate barrier crossing. By frequently swapping the replicas and the original system during the MD, the simulations "travel" on different free energy surfaces and easily visit different conformational zones. Finally, only the samples in the original system (with unmodified free energy surface) are collected. The replicas are artificial and are only used to overcome the energy barriers. REST2 modifies the free energy surfaces by scaling (reducing) the force constants of the "solute" molecules in the simulation system. In this case, the protein and the ligands were considered as "solute"–the force constants of their van der Waals, electrostatic and dihedral terms were subject to scaling–in order to facilitate their conformational changes. The effective temperatures used here for generating the REST2 scaling factors ranged from 310 K to 1000 K for 64 replicas, following a distribution calculated with the Patriksson-van der Spoel approach.[16] Exchange between the replicas was attempted every 1000 simulation steps. This setup resulted in an average exchange probability of ~30 % throughout the simulation course.

*Setup of MD simulations*

The initial models of AVP V2R in inactive state were built using Modeller v9.15 based on the cryo-EM structures of AVP-V2R-Gs (PDBs 7DW9 [17] and 7BB6 [1]) and the X-ray crystal structure of the oxytocin receptor in inactive state (PDB 6TPK[18]). In this model, the intracellular half of V2R was based on the oxytocin receptor inactive state. The *apo,* wt and mutant forms were generated using the same procedure, excluding AVP. The MCF14 and TVP bound forms were obtained by docking the ligands to the *apo* forms, using Autodock Vina.[19] A grid box was set to encompass the pocket with a 0.375 Å grid point spacing. Ligands and pocket residues were set flexible during docking. Paramagnetic tags were added to the ligand-bound forms above. The linker to the paramagnetic tag in $^{par}$MCF14 was generated in an arbitrary initial conformation.[19] PACKMOL-Memgen[20] was used to assign the side-chain protonation states and embed the models in a lipid bilayer of POPC and cholesterol in 3:1 ratio. The systems were solvated in a periodic 78 × 78 × 112 $Å^3$ box of explicit water and neutralized with 0.15 M of $Na^+$ and $Cl^-$ ions. We used the Amber ff14SB,[21] GAFF[22] and lipid14[23] force fields, and the TIP3P water models[24] and the Joung-Cheatham ion parameters[25]. Effective point charges of the ligands were obtained by RESP fitting[26] of the electrostatic potentials calculated with the HF/6-31G* basis set. After energy minimization, all-atom MD simulations were carried out using Gromacs 2020[27] patched with the PLUMED 2.3 plugin.[28] Each system was gradually heated to 310 K and pre-equilibrated during 10 ns of brute-force MD in the *NPT*-ensemble. The replica exchange with solute scaling (REST2)[15] technique was used to enhance the MD sampling (see Supplementary Methods for details and Figure S12). We performed 60 ns × 64 replicas of REST2 MD in the *NVT* ensemble for each system. The first 20 ns were discarded for equilibration. The trajectories of the original unmodified replica were collected and analyzed.

#### 2. Supplemental Items

**Table S1.** Ligand binding affinity for wt V2R and variants

| Receptor | $K_d$ Benzazepine-red (nM) | $K_i$ AVP (nM) |
|---|---|---|
| wt V2R | 2.3 ± 0.3[a] | 0.9 ± 0.3 (n=5) |
| $V2R^{K100R}$ | 2.3 ± 1 | 3.2 ± 2.1 |
| $V2R^{K116R}$ | 8.5 ± 1.9 | 3.4 ± 0.8 |
| $V2R^{K268R}$ | 2.9 ± 1.3 | 2.0 ± 1.3 |
| $V2R^{I130N}$ | 6.6 ± 2.5 | 1.4 ± 0.6 |

[a] from ref [1]. Competitive binding experiments were performed as detailed in Materials and Methods, using the benzazepine-red fluorescent ligand as a tracer. Specific binding of the benzazepine-red antagonist is calculated as a FRET ratio (665nm/620nm). $K_d$ and $K_i$ values are mean ± SEM (n = 3, unless otherwise indicated).

**Table S2.** Ligand binding affinity and potency for $V2R^{K100R}$ variant.

| | Binding (Ki nM) | cAMP accumulation (EC50 nM) |
|---|---|---|

| | | |
|---|---|---|
| AVP | 3.2 ± 2.1 | 0.12 ± 0.05 |
| MCF14 | 11.4 ± 0.4 | 26.9 ± 3.3 |
| Tolvaptan | 1.3 ± 0.4 | n.a[a] |

[a] n.a: not applicable

Competitive binding experiments were performed as detailed in Materials and Methods, using the benzazepine-red fluorescent ligand as a tracer. Specific binding of the benzazepine-red antagonist is calculated as a FRET ratio (665nm/620nm). Accumulation of cAMP is calculated as a FRET ratio (665nm/620nm) and measured in the presence of increasing concentrations of ligands (AVP, MCF14, TVP). $K_i$ and EC50 values are mean ± SEM (n = 3).

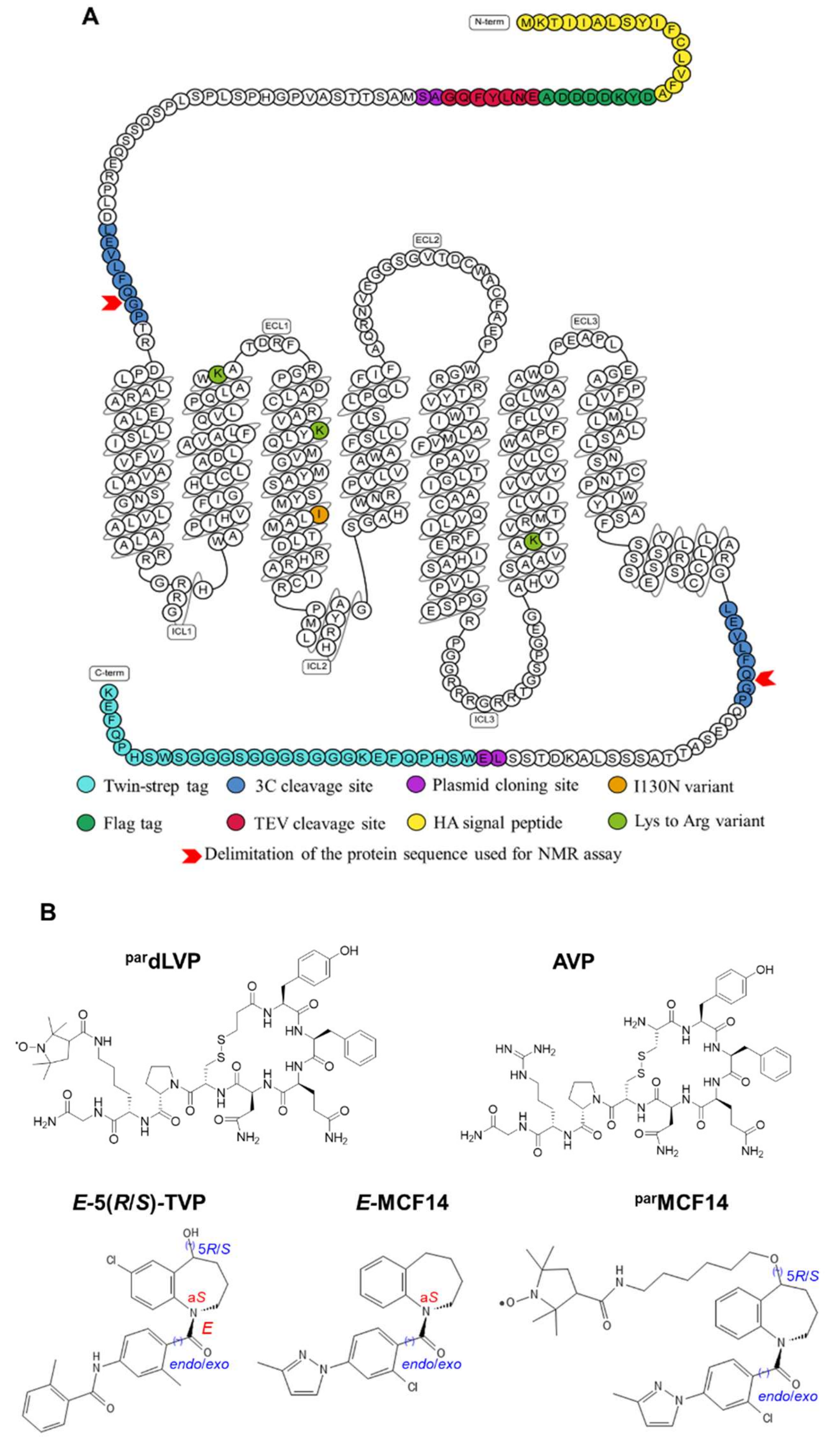


***Figure S1.*** Engineering of V2R and ligands. (A) Snake plot of the V2R sequence used in this study. The hemagglutinin (HA) signal peptide (MKTIIALSYIFCLVFA, in yellow) followed by a Flag-tag (DYKDDDDA, in dark green) and a TEV protease cleavage site (ENLYFQG, in red) were added at the N-terminus, while a Twin-strep-tag® (WSHPQFEKGGGSGGGSGGGSWSHPQFEK, in cyan) was inserted at the C-terminus. In addition, N22 was substituted with a glutamine residue to avoid N-glycosylation, and C358 mutated into an alanine to eliminate potential intermolecular disulfide bridges during solubilization and purification. Two HRV3C protease cleavage sites (LEVLFQGP, in blue) were also added to remove the N- and C-termini and enhance the NMR spectra quality, one in the N-terminus between D30 and T31 and one in the C-terminus between G345 and Q354 (replacing R346-TPPSLG-P353), indicated by two red arrows. M1 and L2 residues after truncation were replaced by A and S residues (purple), and LE residues (purple) were added just before the Twin-strep-tag®, during subcloning (by introduction of a Nhe1 and a Xho1 restriction site, respectively). Mutation sites of K100$^{2.65}$R, K116$^{3.29}$R, and K268$^{6.32}$R are indicated in light green, and I130$^{3.43}$N in orange. Arrows indicate N- and C-termini of the V2R protein after HRV3C cleavage. (B) Chemical structures of the ligands (see Figure S4 for the synthesis).. For TVP and MCF14, both *endo/exo* isomers were tested in MD simulations. The *endo*-isomer turned out to bind more stably. Therefore, only the *endo*-isomer of parMCF14 was used for MD simulations.

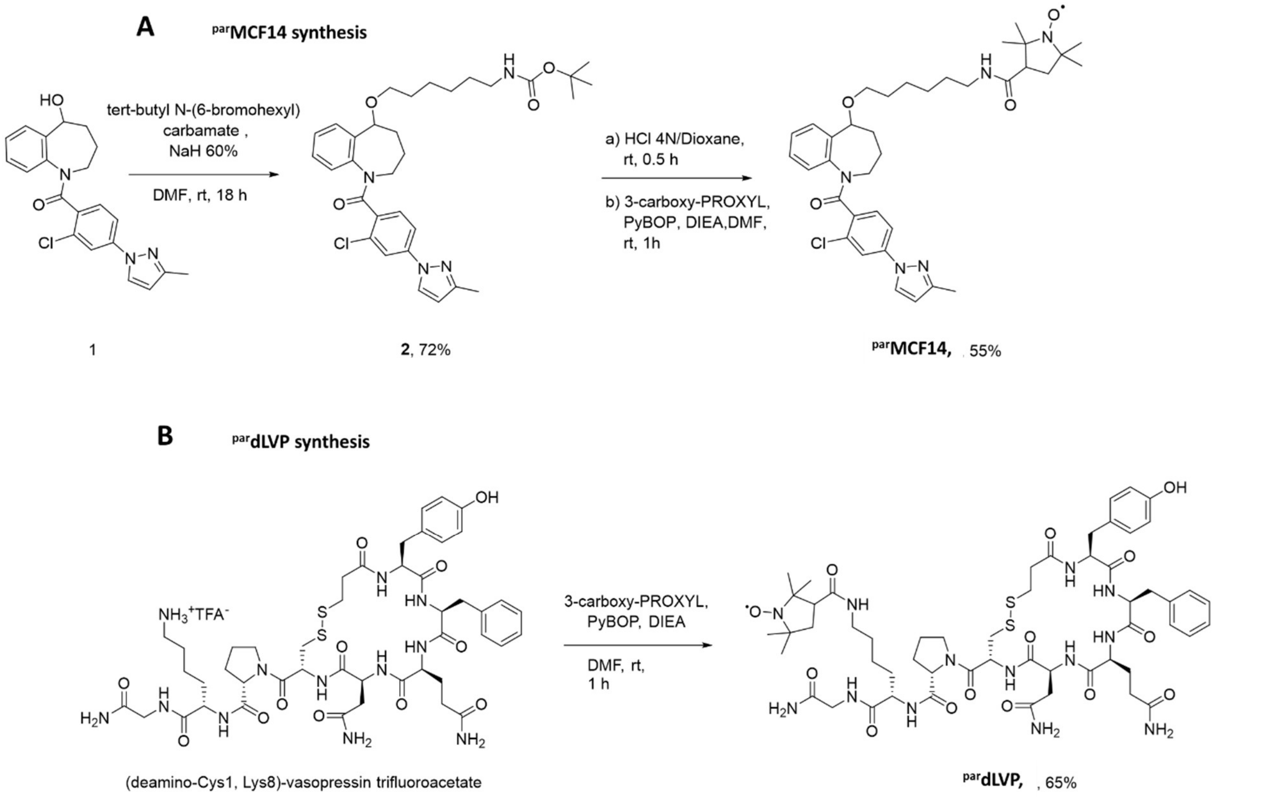


***Scheme S1*.** Synthesis of paramagnetic ligands. (A) $^{par}$MCF14 and (B) $^{par}$dLVP. The procedures are fully detailed in the supplementary methods section.

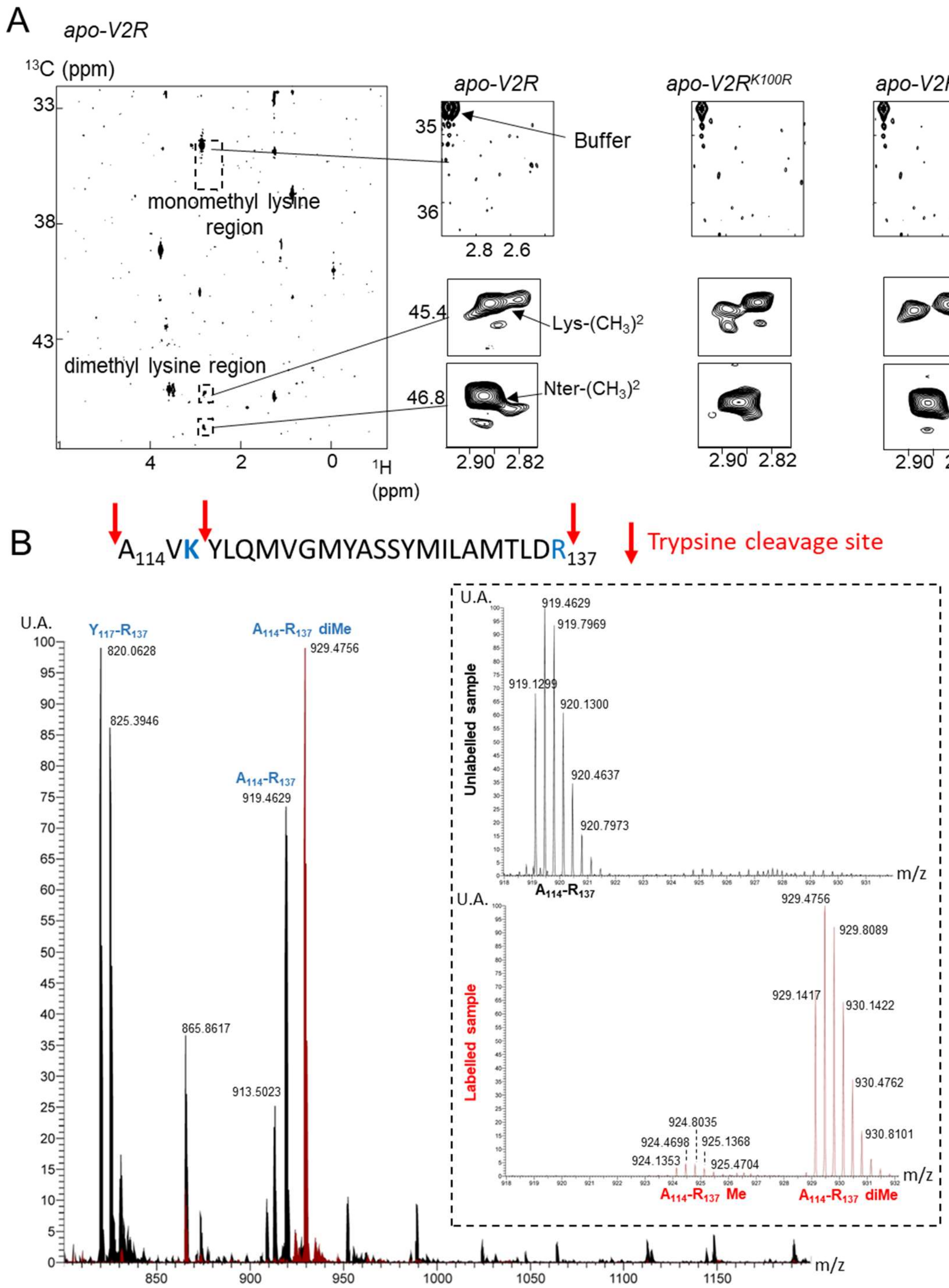


C

| | Sequence | Peak area (10e6) Unlabelled | Peak area (10e6) Labelled - Me | Peak area (10e6) Labelled - diMe | Ratio (Unlabelled : Me : diMe) (%) |
|---|---|---|---|---|---|
| Unlabelled | A114 - R137 | 1,440 | / | 18 | 98.8 : 0 : 1.2 |
| | Y117 - R137 | 3,785 | / | / | 100 : 0 : 0 |
| Labelled | A114 - R137 | 1 | 63 | 1,476 | 0.1 : 4.1 : 95.8 |
| | Y117 - R137 | 0 | / | / | 0 : 0 : 0 |

***Figure S2.*** Analysis of lysine methylation by NMR and mass spectrometry. (A) Identification of the mono-lysine and dimethyl-lysine regions in a representative HMQC spectrum acquired from purified samples of wild-type and mutant apo-V2R. The black boxes and their corresponding close-up views indicate location of monomethyl and dimethyl regions. The zoom for the monomethyl region was centered on the theoretical region 2,4-3 ppm/34.5-36.5 ppm in the $^{1}H$ and $^{13}C$ dimension respectively. The signal seen in the monomethyl lysine region only arises from the Hepes buffer, no specific signal is detected for monomethylated lysines. In the dimethyl-lysine region, peaks arising from labeled lysines (Lys-$(CH_3)^2$) and the N-terminal of the V2R (Nter-$(CH_3)^2$, generated after HRV3C cleavage) are shown. (B) Comparison of unlabeled and $^{13}C$-methylated V2R by mass spectrometry (see Materials and Methods section). The analysis has been performed on the V2R$^{K100R}$ mutant, displaying only 2 lysine residues after HRC3C cleavage (K116 and K268, see Supplementary Figure S1A). Only the tryptic peptide containing K116 is represented in the figure (A114-R137, the trypsine cleavage sites are shown by red arrows). Equivalent results were obtained for the peptide encompassing K268 (R252-R271). Sum of MS spectra from 67.3 to 67.8 min (m/z limited to 800-1200) obtained from unlabeled and $^{13}C$-methylated V2R are superimposed: unlabeled peptides are shown in black, whereas the $^{13}C$-methylated peptides are depicted in red. Peaks arising from A114-R137 and Y117-R137 can be detected in the unlabeled sample. Only a major peak corresponding to $^{13}$C-dimethylated lysine A114-R137 peptide is seen in the labeled sample. Close-up views are displayed on the right: in the unlabeled sample, only signal for A114-R137 is measured, whereas in the labeled sample, monomethylated A114-R137 (very low intensity) and dimethylated A114-R137 (high intensity) are measured. A m/z difference of 10 (919 versus 929) between unlabeled and dimethyl-labeled lysine is measured, which means a mass difference of 30 (the peptide contains 3 charges) corresponding to two methyl groups. In addition, a m/z difference of 5 (919 versus 924) between unlabeled and monomethyl-labeled lysine is measured, which means a mass difference of 15 (3 charges) corresponding to one methyl group. (C) Methylation ratio of the representative A114-R137 tryptic peptide of the V2R$^{K100R}$ mutant. Peak areas corresponding to the unlabeled, mono- and dimethyl-labeled lysine are calculated. In the unlabeled sample, A114-R137 and Y117-R137 peptides are detected: for both peptides, the unlabeled species are major (1,440 and 3,785 units, respectively), as compared to no signal (/) or very minor signal (18 units) for the dimethyl-labeled lysine in the A114-R137 peptide. In the labeled sample, the A114-R137 dimethyl-labeled lysine peptide is almost exclusive (1,476 units versus 1 for unlabeled and 63 for the monomethyl-labeled), and the Y117-R137 is not detected (a 0 value means <1). Therefore, it appears that reductive dimethylation of the V2R onto its lysine residues is nearly complete with more than 95% efficiency.

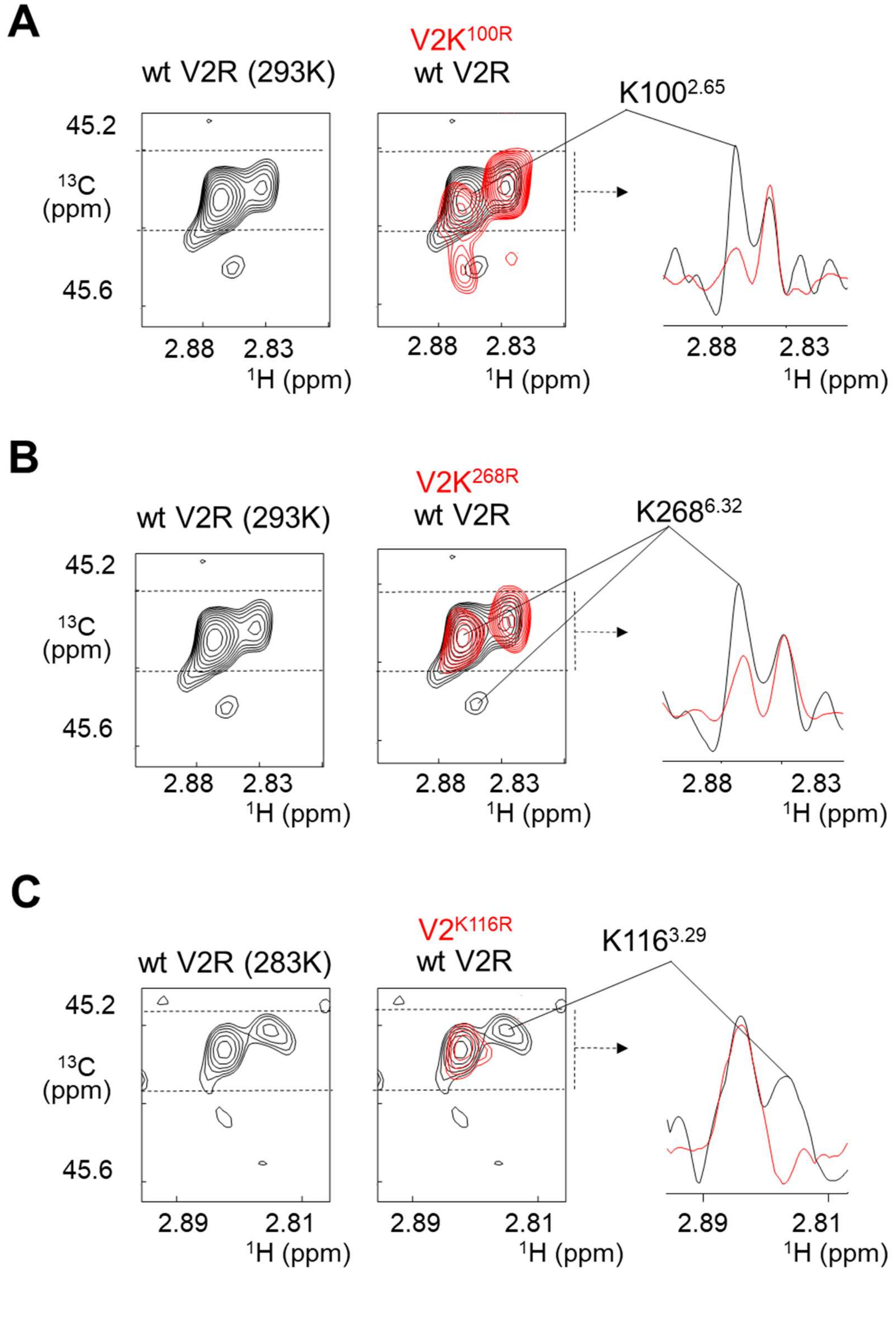

***Figure S3.*** Assignment of correlation peaks of dimethylated lysines. Extracted HMQC spectrum of native V2R (black) is superimposed on that of the mutants (red): **(A)** K100$^{2.65}$R, **(B)** K268$^{6.32}$R and **(C)** K116$^{3.29}$R. Experiments were run at 293 K, except for the assignment of the K116 resonance (C). On the right panels, we highlight the peak disappearance for each mutant (red line) as compared to native V2R (black line) in the $^{1}$H dimension. The 1D projections along the $^{13}$C dimension of $^{1}$H rows between the two dashed lines are indicated by an arrow (right panels).

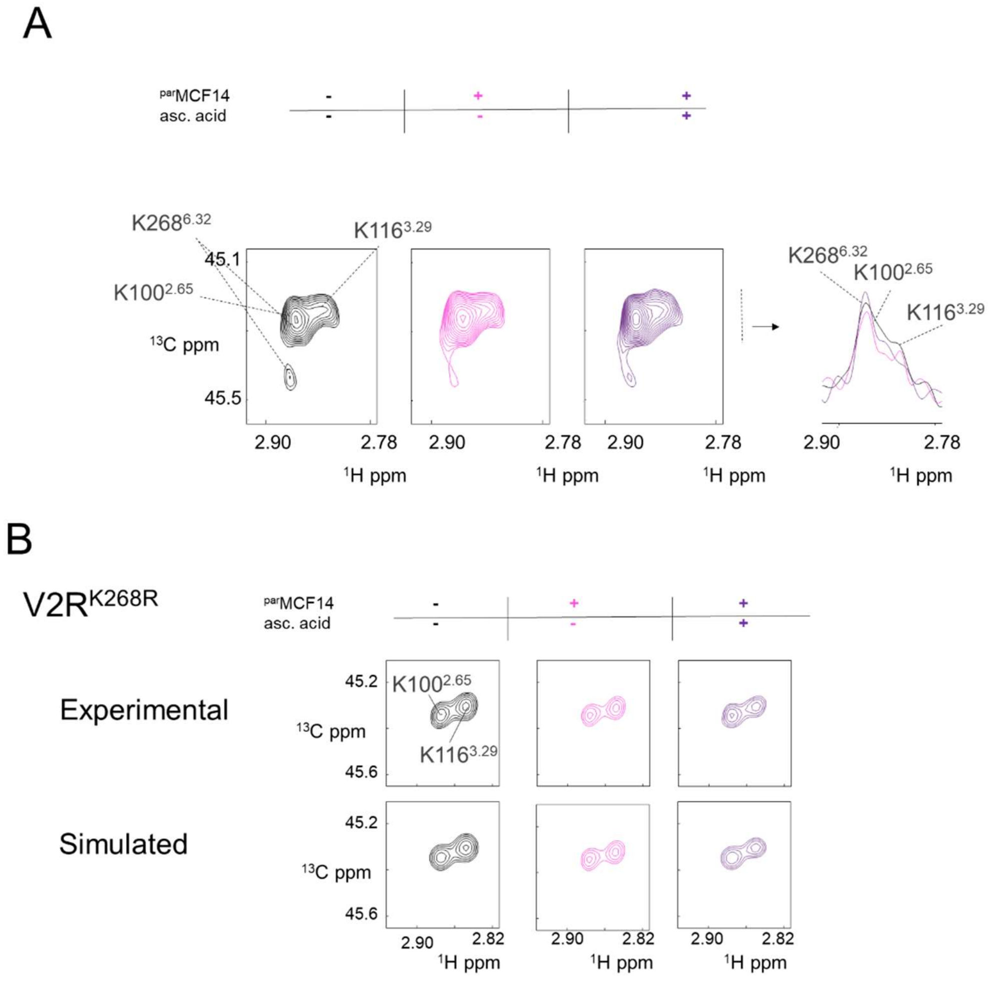


***Figure S4.*** Effect of $^{par}$MCF14 binding and reduction by ascorbic acid on the lysine HMQC correlation peaks of wt V2R and fitting procedure. (A) The 1D projection along the $^{13}$C dimension of $^{1}$H rows is indicated by an arrow, showing a specific effect according to each lysine. (**B**) Experimental and simulated HMQC correlation of V2R$^{K268R}$ in the *apo*-, $^{par}$MCF- and $^{par}$MCF14$_{red}$-bound states**.**

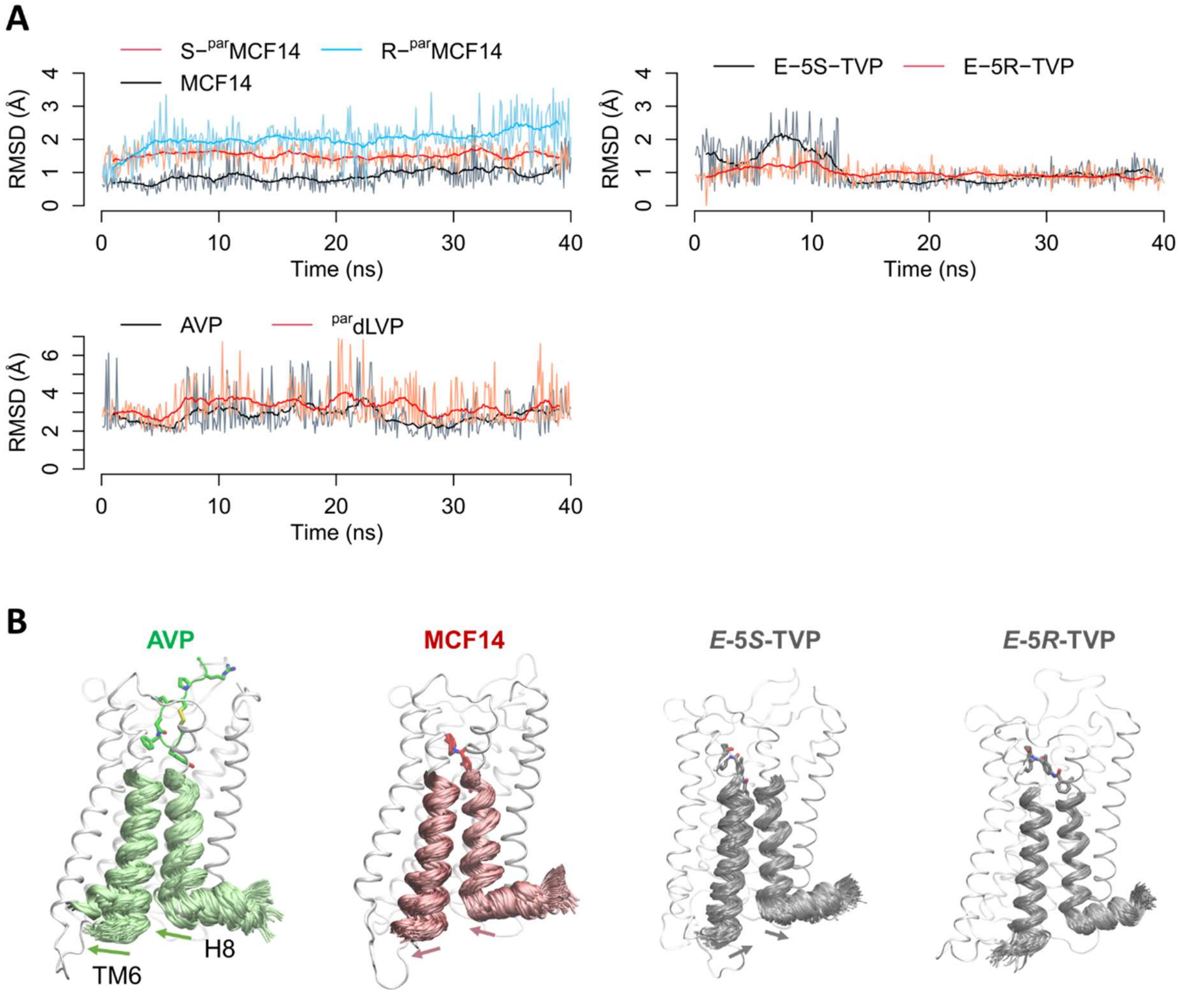


***Figure S5.*** Conformational dynamics of V2R and ligands during MD simulations. (A) Ligand RMSD calculated for the heavy atoms of MCF14 and TVP, or the backbone atoms of AVP and $^{par}$dLVP, excluding the paramagnetic tag. (B) Movements of TM6, TM7 and H8 on the intracellular side of V2R in the simulations.

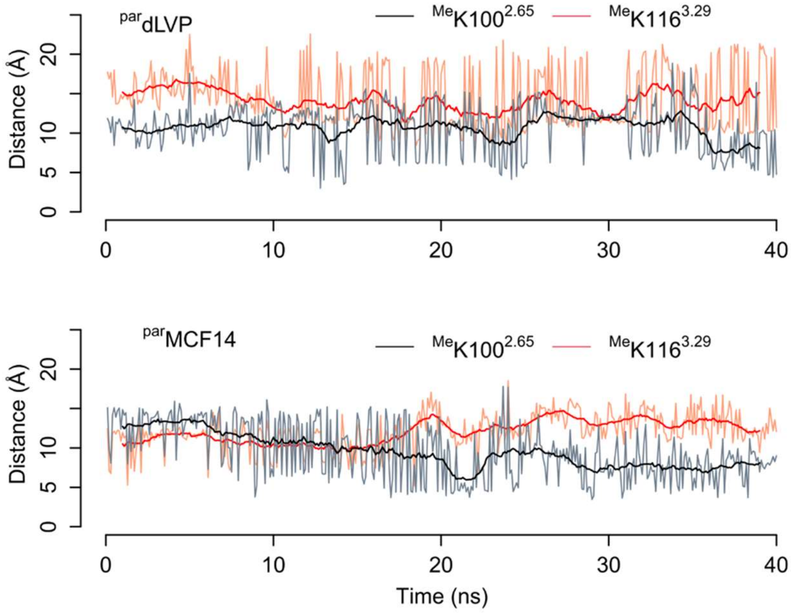

***Figure S6.*** Distances between the paramagnetic tag and dimethylated K100$^{2.65}$ and K116$^{3.29}$ during the MD simulations. The distance is measured between the nitroxide oxygen atom and nitrogen atom of the lysine side chain.

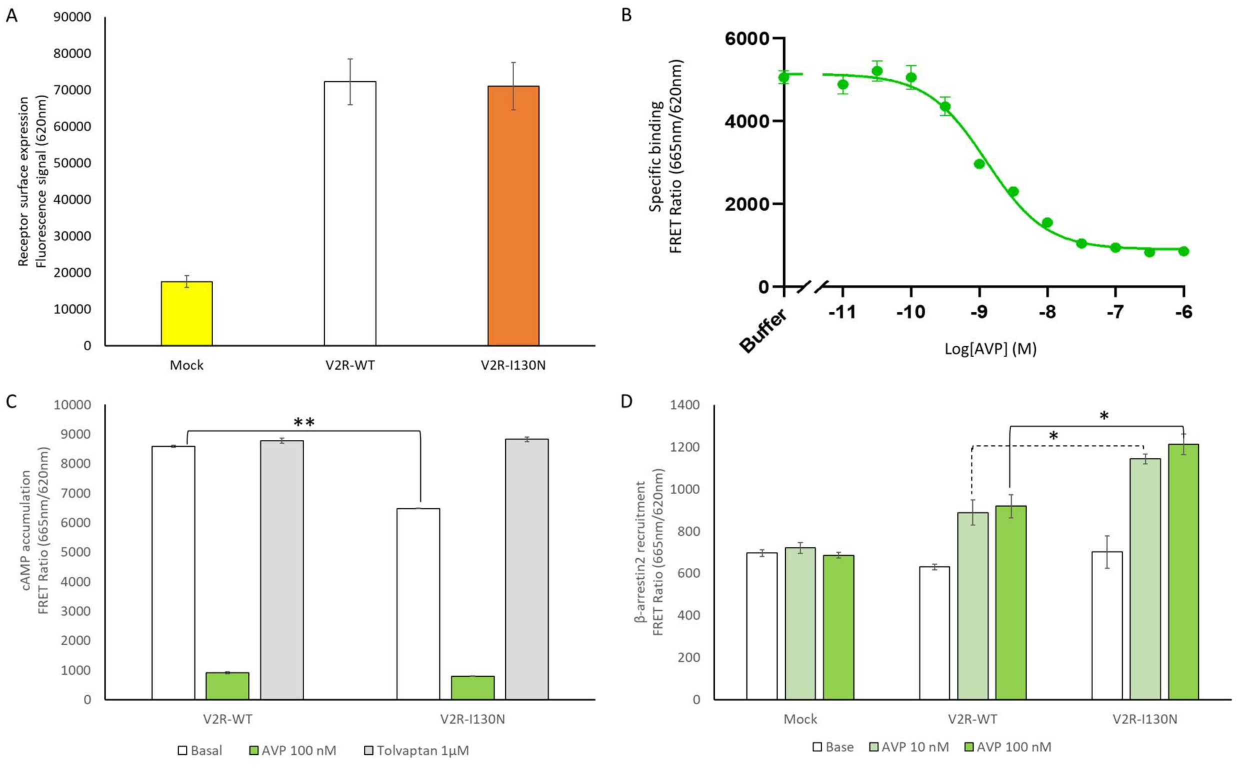


***Figure S7.*** Pharmacological and functional properties of the mutant V2R$^{I130N}$. (A) Evaluation of the receptor expression level at the cell surface by measuring the fluorescence signal at 620 nm (see Materials and Methods for details). The expression of the I130N mutant is compared to that of wt V2R. (B) Binding of AVP to the mutant V2R$^{I130N}$ is illustrated as the FRET ratio (665nm/620nm x 10,000). Specific binding of the fluorescent benzazepine-red antagonist is shown. For each competition curve, the tracer was used at 5 nM with or without increasing concentrations of AVP. (C) Basal and ligand-induced V2R-dependent cAMP accumulation measured by the FRET ratio (665nm/620nm x 10,000). The functional properties of the mutant V2R$^{I130N}$ are compared to those of wt V2R. The cAMP accumulation is compared to that measured in the presence of 100 nM of AVP (in green) or 1 µM of TVP (in grey). (D) V2R-dependent recruitment of β-arrestin2 to AP2 measured by the FRET ratio (665nm/620nm x 10,000) in the presence of 0 nM (basal), 10 nM or 100 nM of AVP. The capacity of β-arrestin2 recruitment of the mutant V2R$^{I130N}$ is compared to that of wt V2R. Data are mean ± SEM of 3 technical replicates where statistical significance is assessed by T-test (*, $p < 0.05$; **, $p < 0.01$).

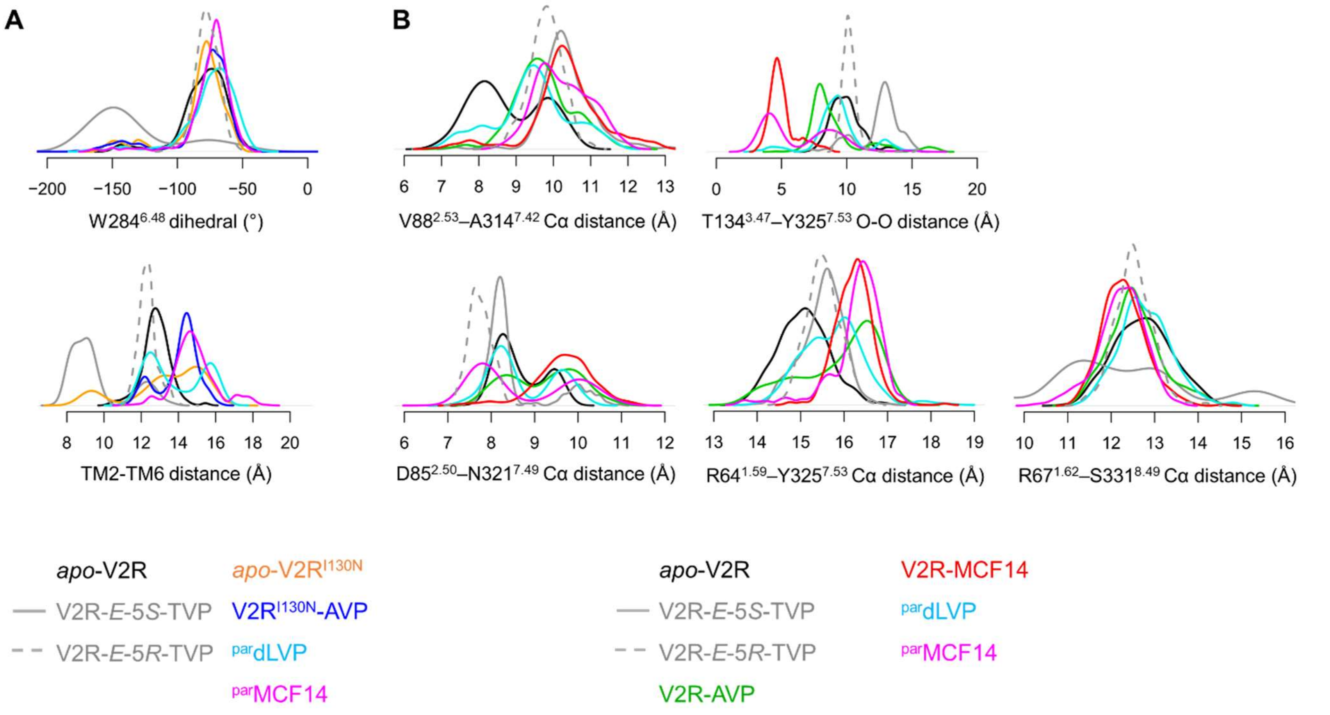


***Figure S8.*** Probability density distribution of the conformational features measured during the MD simulations**. (A)** and **(B)** are supplementary to Figure 3 and Figures **4** and **5**, respectively.

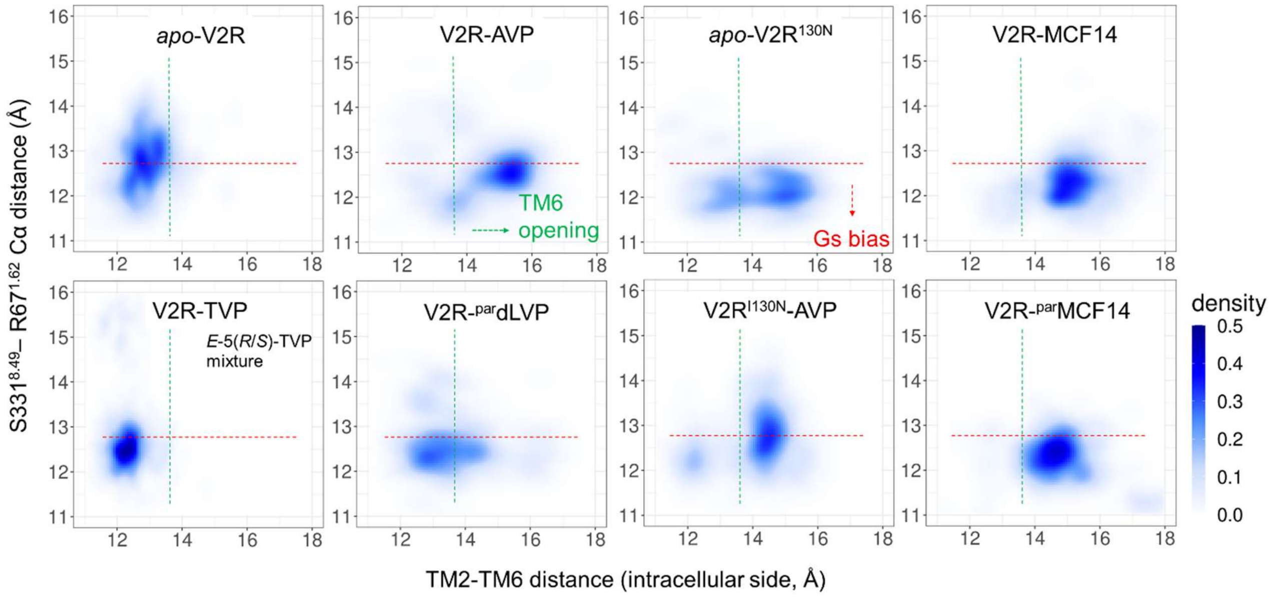


***Figure S9.*** Probability density map of two conformational features during the MD simulations. TM6 opening (x-axis) is considered as an indicator of the canonical activation, whereas H8-TM1 closure (y-axis) is used as an indicator of the G-protein bias. TM6 opening is measured by the center-of-mass distance between the backbones of I74$^{2.39}$-F77$^{2.42}$ and V266$^{6.30}$-T269$^{6.33}$. H8-TM1 closure is measured by the Cα distance between S331$^{8.49}$ and R67$^{1.62}$.

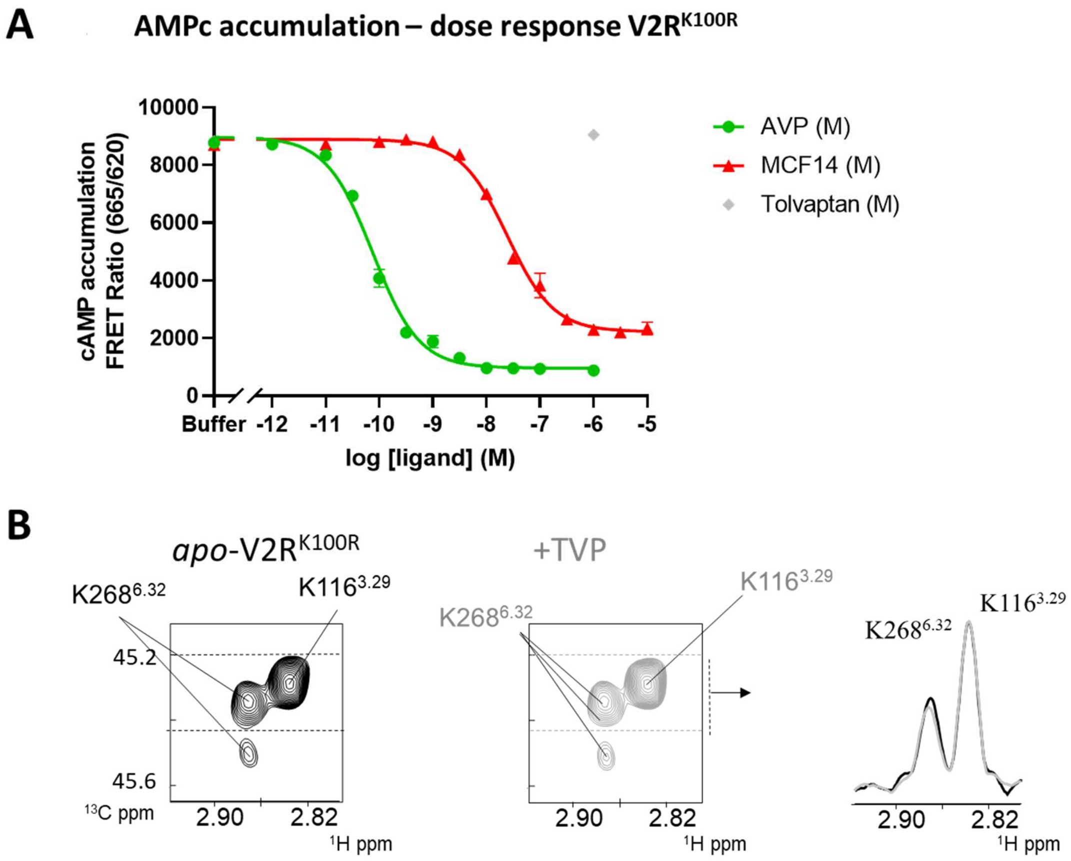


***Figure S10.*** Functional properties of V2R$^{K100R}$. (A) Ligand dose-response curves of V2R-dependent cAMP accumulation measured by FRET ratio (665nm/620nm x 10,000). The cAMP accumulation was quantified using the cAMP Dynamic 2 kit (PerkinElmer CisBio) as indicated in Materials and Methods and is shown in the presence of increasing concentrations of AVP (green), MCF14 (red), and TVP (in grey). Data are mean ± SEM of 3 technical replicates. **(B)** Extracted HMQC spectra of V2R$^{K100R}$ in *apo* form (black) and in the presence of saturating concentrations of the TVP antagonist (grey). 1D spectra on the right correspond to 1D projections along the $^{13}$C dimension of $^{1}$H rows between the two dashed lines. The spectra of the *apo* and TPV-bound states are almost identical, indicating that the *apo* form is mainly inactive.

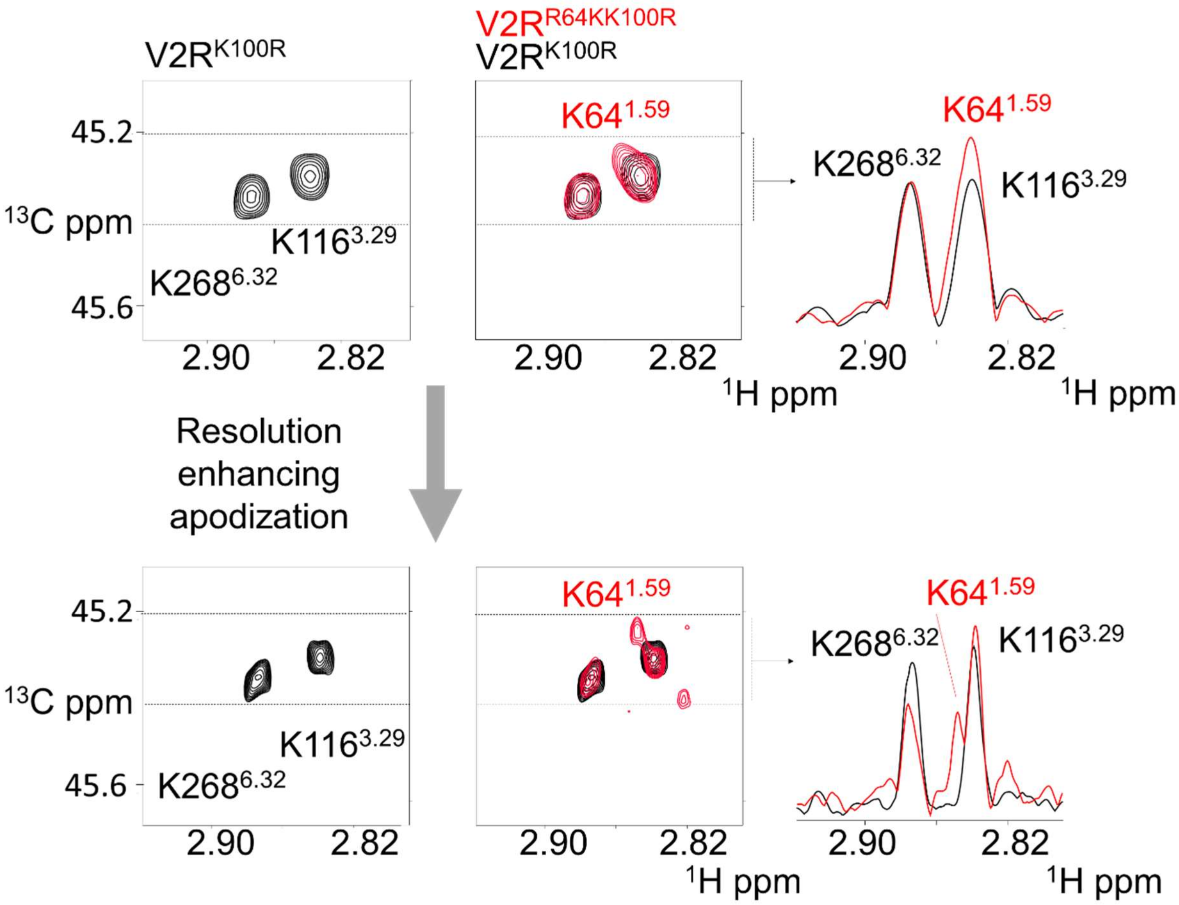


***Figure S11.*** Assignment of correlation peak of the probe K64 of V2R64K-K100R. Extracted HMQC spectra of V2$^{K100R}$ (black) and V2$^{R64K-K100R}$ (red). The bottom spectra are processed with a sinus function. 1D spectra on the right correspond to 1D projections along the $^{13}$C dimension of $^{1}$H rows between the two dashed lines.

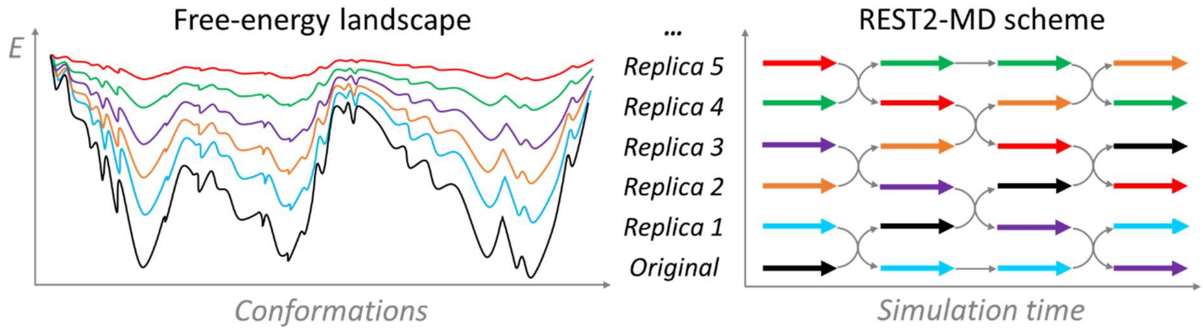


***Figure S12.*** Scheme of REST2 protocol. REST2 is a type of Hamiltonian replica exchange simulation scheme, which performs many replicas of the same MD simulation system simultaneously. The replicas have modified free energy surfaces, in which the barriers are easier to cross than in the original system. By frequently swapping the conformations between neighbor replicas during the MD, the simulations "travel" on different free energy surfaces and easily visit different conformational zones. Finally, only the samples on the original free energy surface are collected. REST2, in particular, modifies the free energy surfaces by reducing the energy barriers of conformational changes of "solute" molecules in the simulation system. Here, the protein and the ligands were considered as "solute", in order to facilitate their conformational changes.

**Supplementary references**


[1] J. Bous, H. Orcel, N. Floquet, C. Leyrat, J. Lai-Kee-Him, G. Gaibelet, A. Ancelin, J. Saint-Paul, S. Trapani, M. Louet, R. Sounier, H. Déméné, S. Granier, P. Bron, B. Mouillac, *Sci. Adv.* **2021**, *21*, eabg5628.

[2] F. Jean-Alphonse, S. Perkovska, M.-C. C. Frantz, T. Durroux, C. Mejean, D. Morin, S. Loison, D. Bonnet, M. Hibert, B. Mouillac, C. Mendre, *J Am Soc Nephrol* **2009**, *20*, 2190–2203.

[3] J. P. Morello, M. Bouvier, U. E. Petäjä-Repo, D. G. Bichet, *Trends Pharmacol. Sci.* **2000**, *21*, 466–469.

[4] M. P. Bokoch, Y. Zou, S. G. F. Rasmussen, C. W. Liu, R. Nygaard, D. M. Rosenbaum, J. J. Fung, H. J. Choi, F. S. Thian, T. S. Kobilka, J. D. Puglisi, W. I. Weis, L. Pardo, R. S. Prosser, L. Mueller, B. K. Kobilka, *Nature* **2010**, *463*, 108–112.

[5] R. Sounier, C. Mas, J. Steyaert, T. Laeremans, A. Manglik, W. Huang, B. K. Kobilka, H. Déméné, S. Granier, *Nature* **2015**, *524*, 375–378.

[6] J. Bous, A. Fouillen, H. Orcel, S. Trapani, X. Cong, S. Fontanel, J. Saint-Paul, J. Lai-Kee-Him, S. Urbach, N. Sibille, R. Sounier, S. Granier, B. Mouillac, P. Bron, *Sci. Adv.* **2022**, *10.1126/sc*, DOI 10.1101/2022.02.11.480047.

[7] J. Cox, M. Mann, *Nat. Biotechnol.* **2008**, *26*, 1367–1372.

[8] B. Schilling, M. J. Rardin, B. X. MacLean, A. M. Zawadzka, B. E. Frewen, M. P. Cusack, D. J. Sorensen, M. S. Bereman, E. Jing, C. C. Wu, E. Verdin, C. R. Kahn, M. J. MacCoss, B. W. Gibson, *Mol. Cell. Proteomics* **2012**, *11*, 202–214.

[9] F. Delaglio, S. Grzesiek, G. W. Vuister, G. Zhu, J. Pfeifer, A. Bax, *J. Biomol. NMR 1995 63* **1995**, *6*, 277–293.

[10] B. A. Johnson, R. A. Blevins, *J. Biomol. NMR 1994 45* **1994**, *4*, 603–614.

[11] S. Loison, M. Cottet, H. Orcel, H. Adihou, R. Rahmeh, L. Lamarque, E. Trinquet, E. Kellenberger, M. Hibert, T. Durroux, B. Mouillac, D. Bonnet, *J. Med. Chem.* **2012**, *55*, 8588–8602.

[12] J. M. Zwier, T. Roux, M. Cottet, T. Durroux, S. Douzon, S. Bdioui, N. Gregor, E. Bourrier, N. Oueslati, L. Nicolas, N. Tinel, C. Boisseau, P. Yverneau, F. Charrier-Savournin, M. Fink, E. Trinquet, *J. Biomol. Screen.* **2010**, *15*, 1248–1259.

[13] J. Tenenbaum, M. A. Ayoub, S. Perkovska, A. L. Adra-Delenne, C. Mendre, B. Ranchin, G. Bricca, G. Geelen, B. Mouillac, T. Durroux, D. Morin, *PLoS One* **2009**, *4*, e8383.

[14] M. C. Frantz, L. P. Pellissier, E. Pflimlin, S. Loison, J. Gandia, C. Marsol, T. Durroux, B. Mouillac, J. A. J. Becker, J. Le Merrer, C. Valencia, P. Villa, D. Bonnet, M. Hibert, *J Med Chem* **2018**, *61*, 8670–92.

[15] L. Wang, R. A. Friesner, B. J. Berne, *J. Phys. Chem. B* **2011**, *115*, 9431–9438.

[16] A. Patriksson, D. Van Der Spoel, *Phys. Chem. Chem. Phys.* **2008**, *10*, 2073–2077.

[17] F. Zhou, C. Ye, X. Ma, W. Yin, T. I. Croll, Q. Zhou, X. He, X. Zhang, D. Yang, P. Wang, H. E. Xu, M. W. Wang, Y. Jiang, *Cell Res.* **2021**, *31*, 929–931.

[18] Y. Waltenspühl, J. Schöppe, J. Ehrenmann, L. Kummer, A. Plückthun, *Sci. Adv.* **2020**, *6*, 1–12.

[19] J. Eberhardt, D. Santos-Martins, A. F. Tillack, S. Forli, *J. Chem. Inf. Model.* **2021**, *61*, 3891–3898.

[20] S. Schott-Verdugo, H. Gohlke, *J. Chem. Inf. Model.* **2019**, *59*, DOI 10.1021/ACS.JCIM.9B00269.

[21] K. Lindorff-Larsen, S. Piana, K. Palmo, P. Maragakis, J. L. Klepeis, R. O. Dror, D. E. Shaw, *Proteins* **2010**, *78*, 1950–1958.

[22] J. Wang, R. M. Wolf, J. W. Caldwell, P. A. Kollman, D. A. Case, *J. Comput. Chem.* **2004**, *25*, 1157–1174.

[23] C. J. Dickson, B. D. Madej, Å. A. Skjevik, R. M. Betz, K. Teigen, I. R. Gould, R. C. Walker, *J. Chem. Theory Comput.* **2014**, *10*, 865–879.

[24] W. L. Jorgensen, J. Chandrasekhar, J. D. Madura, R. W. Impey, M. L. Klein, *J. Chem. Phys.* **1998**, *79*, 926.

[25] I. S. Joung, T. E. Cheatham, *J. Phys. Chem. B* **2008**, *112*, 9020–9041.

[26] J. Wang, P. Cieplak, P. Kollman, *J. Computational Chem.* **2000**, *21*, 12.

[27] D. Van Der Spoel, E. Lindahl, B. Hess, G. Groenhof, A. E. Mark, H. J. C. Berendsen, *J. Comput. Chem.* **2005**, *26*, 1701–1718.

[28] G. Tribello, M. Bonomi, D. Branduardi, C. Bussi, *Comput. Phys. Commun.* **2014**, *185*, 604–13.
